# Influenza A M2 recruits M1 to the plasma membrane: a fluorescence fluctuation microscopy study

**DOI:** 10.1101/2021.05.06.442926

**Authors:** Annett Petrich, Valentin Dunsing, Sara Bobone, Salvatore Chiantia

## Abstract

Influenza A virus (IAV) is a respiratory pathogen that causes seasonal epidemics with significant mortality. One of the most abundant proteins in IAV particles is the matrix protein 1 (M1), which is essential for the virus structural stability. M1 organizes virion assembly and budding at the plasma membrane (PM), where it interacts with other viral components. The recruitment of M1 to the PM as well as its interaction with the other viral envelope proteins (hemagglutinin (HA), neuraminidase, matrix protein 2 (M2)) is controversially discussed in previous studies. Therefore, we used fluorescence fluctuation microscopy techniques (i.e., scanning fluorescence cross-correlation spectroscopy and Number and Brightness) to quantify the oligomeric state of M1 and its interactions with other viral proteins in co-transfected as well as infected cells. Our results indicate that M1 is recruited to the PM by M2, as a consequence of the strong interaction between the two proteins. In contrast, only a weak interaction between M1 and HA was observed. M1-HA interaction occurred only in the case that M1 was already bound to the PM. We therefore conclude that M2 initiates the assembly of IAV by recruiting M1 to the PM, possibly allowing its further interaction with other viral proteins.

**Statement of Significance:** Influenza A virus (IAV) is a pathogen responsible for epidemics and occasional pandemics and, therefore, a significant burden on health systems. To develop innovative therapeutic approaches, a deeper understanding of the viral replication cycle is needed. For example, during the formation of new virions in infected cells, several viral components must assemble at the plasma membrane, but the molecular interactions involved in this process are not clearly understood. In this work, we use quantitative fluorescence microscopy methods to monitor the interplay between several viral proteins in live cell models. Our results underline the importance of the interactions between two specific proteins (M1 and M2) and shed light on the first steps in IAV assembly.

## Introduction

Influenza A viruses (IAVs) belong to the family of the *Orthomyxoviridae*. These pathogens represent a substantial global health burden, being associated with significant morbidity and mortality through frequent epidemics and several pandemics (1, 2). IAV is enveloped by a lipid bilayer that is derived from the host cell membrane and contains two integral transmembrane glycoproteins (i.e. hemagglutinin (HA) and neuraminidase (NA)) and one transmembrane protein with a proton-selective ion channel activity (i.e. the matrix protein 2 (M2)) (3, 4). The envelope protein HA is a homotrimeric type I transmembrane glycoprotein and is the major surface protein of IAV particles (5-7). HA plays a major role in viral entry by mediating the attachment of the virus to cell surface sialic acid molecules, membrane fusion after internalization, and the release of viral genome into target cells (5-8). The surface protein NA is a homotetrameric type II transmembrane glycoprotein that facilitates the release of newly synthesized virus particles from the infected cells by enzymatic cleavage of the cell surface receptor molecules (5-8). Additionally, a small amount of homotetrameric M2 molecules are embedded in the viral envelope (approximately 16 to 20 molecules in a virus, compared to ca. 300-400 HA and 50 NA copies) (6, 7). M2 is a type III transmembrane protein which functions as proton channel activated by acidic pH and is important for genome unpacking during virus entry (7-9). Moreover, it was shown that M2 is connected to virus morphology, production of infectious virus particles, and membrane scission (9-13). All the three envelope proteins are transported from the trans-Golgi network to the apical plasma membrane via the secretory pathway (8, 9, 14). Both glycoproteins, HA and NA, are supposed to be enriched in lipid “raft” microdomains at the virion budding site, whereas M2 was suggested to localize to the edges of such domains (8, 14-16).

The luminal side of the viral envelope is coated with the matrix protein 1 (M1), which forms the viral nucleocapsid in close contact to the lipid membrane (17-20), binds the viral ribonucleoproteins (vRNPs) (4, 21), and is supposed to interact with viral surface proteins (10, 11, 22-24). Moreover, M1 is the most abundant, highly conserved protein in IAV particles and is important for several processes during viral replication, including the regulation of capsid disassembly, virus budding and morphogenesis (3, 8, 25). Interestingly, M1 lacks an apical transport signal, implying that the membrane localization of M1 in infected cells might be due to piggyback transport with HA, NA, M2 or vRNPs (26, 27). For this reason, various hypotheses regarding the association of M1 to the plasma membrane (PM) have been proposed over the years. First, several studies established that M1 associates with negatively charged lipids in model membranes (17-20, 28, 29). Nevertheless, such interactions appear not to be sufficient for the actual association of M1 to the PM in non-infected cells (i.e. in cells expressing M1 as the only viral protein) (17, 27). Accordingly, M1 was proposed to interact with the cytoplasmic tails of HA and NA during their apical transport (22-24, 30, 31), as well as with the cytoplasmic tails of M2 at the assembly site (10, 11, 27). Interactions between M1 and HA, NA, or M2 have been investigated via bulk biochemistry methods (e.g. by altered detergent solubility (22, 24), increased membrane association (31) of M1 in the presence of HA or NA, or co-immunoprecipitation of M1 in the presence of M2 (10, 11, 32)). Nevertheless, no clear consensus has been reached regarding the role of HA, NA or M2 in recruiting M1 to the PM and its subsequent incorporation into virions (11, 33-37). In conclusion, the molecular mechanisms involved in M1-driven IAV assembly are not fully understood and the specific interactions between M1 and other viral surface proteins have not yet been quantified directly in living cells.

To obtain quantitative information on how protein-protein interactions (e.g. M1-M1 or M1-HA) occur in the native cellular environment, minimally invasive approaches (e.g. fluorescence fluctuation spectroscopy) are needed (38). Here, we apply (cross-correlation) Number and Brightness (N&B and ccN&B) as well as scanning fluorescence (cross-)correlation spectroscopy (sFCS and sFCCS) analysis in living cells to quantify oligomeric state, concentration and diffusion dynamics of the viral envelope proteins (HA, NA, M2) and M1, as well as their interactions. Our results suggest the presence of a strong interaction between M1 and M2, leading to the recruitment of M1 to the PM in a M2 concentration-dependent manner. We further hypothesize that the interaction between M1 and HA occurs in a subsequent step. Finally, we provide the first experimental evidence of a possible M2 binding-site within the N-terminal domain of M1.

## Materials and Methods

### Plasmids and cloning

The plasmids for the transcription and translation of influenza virus RNAs and proteins of the influenza A/FPV/Rostock/1934 virus (H7N1; FPV) mutant 1 were obtained from Michael Veit (Free University, Berlin, Germany), and previously described (39, 40). The plasmids encoding the fluorescence proteins (FP) mEGFP or mCherry2 linked to a myristoylated and palmitoylated peptide (mp-mEGFP, mp-mCherry2, mp-2x-mEGFP), and the plasmids for cytosolic expression of mEGFP, 2x-mEGFP were previously described (41) and are available on Addgene (Watertown, MA, USA). The plasmids encoding the FP hetero-dimer mCherry2-mEGFP linked to a myristoylated and palmitoylated peptide (mp-mCherry2-mEGFP), and the matrix protein 2 (M2) of FPV with mCherry2 fused to the extracellular terminus of M2 (mCherry2-M2) were previously described (42). Further information regarding other plasmids and constructs used in this work is provided in the Supplemental Information (SI).

### Cell culture and virus infection

Human embryonic kidney (HEK) cells from the 293T line (CRL-3216TM, purchased from ATCC®, Kielpin Lomianki, Poland), and Madin-Darby canine kidney type II (MDCK II) cells (ECACC 00062107, European Collection of Authenticated Cell Cultures, Porton Down, UK) were cultured in phenol red-free Dulbecco’s modified Eagle medium (DMEM) with 10 % fetal bovine serum, 2 mM L-glutamine, 100 U/mL penicillin, and 100 μg/mL streptomycin at 37 °C and 5 % CO_2_. Cells were passaged every 2-3 days when they reached nearly 80 % confluence in tissue culture flask, for no more than 15 times. All solutions, buffers, and media used for cell culture were purchased from PAN-Biotech (Aidenbach, Germany).

For immunostaining experiments, dishes were coated with a 0.01 % (w/v) poly-L-lysine solution (MW 150,000 – 300,000 Da, Sigma-Aldrich, Munich, Germany) before cell seeding.

Information regarding virus propagation, titration and infection is provided in the SI.

### Immunofluorescence

Transfected and infected cells were fixed at the indicated time points with 4 % (w/v) paraformaldeyde (Sigma Aldrich, Taufkirchen, Germany) in DPBS+/+. After 15 min, cells were washed three times with DPBS+/+. Permeabilization was performed with 0.1 % (v/v) Triton X-100® (Sigma Aldrich, Taufkirchen, Germany) for 10 min, and subsequently washed three times with DPBS+/+. Afterwards, cells were incubated with 2 % (w/v) BSA (Sigma Aldrich, Taufkirchen, Germany) in DPBS+/+ for one hour at room temperature. Primary antibody (monoclonal mouse anti-influenza A M2, clone 14C2 (#ab5416, abcam, Cambridge, UK), monoclonal mouse anti-influenza A H7 (#3HI7, HyTest Ltd, Turku, Finland), Clone monoclonal mouse anti-influenza A N1, clone #2F10E12G1 (#AB_2860298, SinoBiological, Eschborn, Germany), monoclonal mouse anti-influenza nucleoprotein, clone A1 (#MAB8257, Millipore trademark of Merck KGaA, Darmstadt,Germany)) were diluted 1:200 or 1:1000 in 0.2 % (w/v) BSA in DPBS+/+, and incubated overnight at 4 °C. After three washing steps with DPBS+/+, cells were incubated with the 1:1000 diluted secondary antibodies (goat anti-mouse AlexaFluor® 488-conjugated or AlexaFluor® 568-conjugated; Thermo Fisher Scientific, Waltham, MA, USA) for one hour at room temperature. Cells were subsequently washed three times with DPBS+/+.

### Confocal microscopy imaging

Microscopy measurements were performed on a Zeiss LSM780 system (Carl Zeiss, Oberkochen, Germany) using a Plan-Apochromat 40×/1.2 Korr DIC M27 water immersion objective and a 32-channel GaAsP detector array. To decrease out-of-focus light, a pinhole with size corresponding to one airy unit (∼39 μm) was used. Samples were excited with a 488-nm argon laser and a 561-nm diode laser. Fluorescence was detected between 499 and 552 nm (mEGFP, AlexaFluor®488) and between 570 and 695 nm (mCherry2), after passing through a MBS 488/561-nm dichroic mirror. For multicolor measurements, fluorophores were excited and detected sequentially for different regions of the spectrum. Confocal imaging was performed with a frame size of 512 × 512 pixels.

Further information regarding setup calibration, sFCCS, (cc)N&B, bi-directional plasmids and multimerization analysis is provided in the SI. A schematic overview of the sFCCS and ccN&B analysis is shown in Figure S1.

### Statistical analysis

Data from at least three independent experiments were pooled and visualized by using GraphPad Prism vs. 9.0.0 (GraphPad Software, LCC, San Diego, CA, USA) or R (R Foundation for Statistical Computing, Vienna, Austria) packages *ggplot2* (43), *ggpubr* (44), and *cowplot* (45). If not otherwise indicated, data were displayed as box plots with single data points corresponding to measurements in single cells. Median values and whiskers ranging from minimum to maximum values are displayed. Quantities in the main text are given as median ± IQR. Sample sizes and p-values are given in each graph and figure captions, respectively. Statistical significance was tested by using D’Agostino-Pearson normality test followed by the one-way ANOVA analysis and the Bonferroni’s multiple comparisons test.

## Code availability

MATLAB custom-written code is available from the corresponding author upon reasonable request.

## Data availability

The datasets analyzed during the current study are available from the corresponding author upon reasonable request.

## Results

### M1 is recruited to the PM by M2 but not by HA or NA

Previous studies have shown that the intracellular localization of the Influenza A matrix protein M1 varies between transfected and infected cells (15, 27). As a starting point for our investigations, we have therefore characterized the behavior of a M1-mEGFP fluorescent construct derived from the avian IAV strain FPV directly in living HEK293T cells. Protein localization was monitored via confocal microscopy either i) when expressed by itself, ii) in the presence of all other viral proteins (i.e., via the reverse genetic plasmid system and unlabeled M2 termed here as “all”), or iii) in FPV infected cells (Figure 1 A, B).

**Figure 1:**
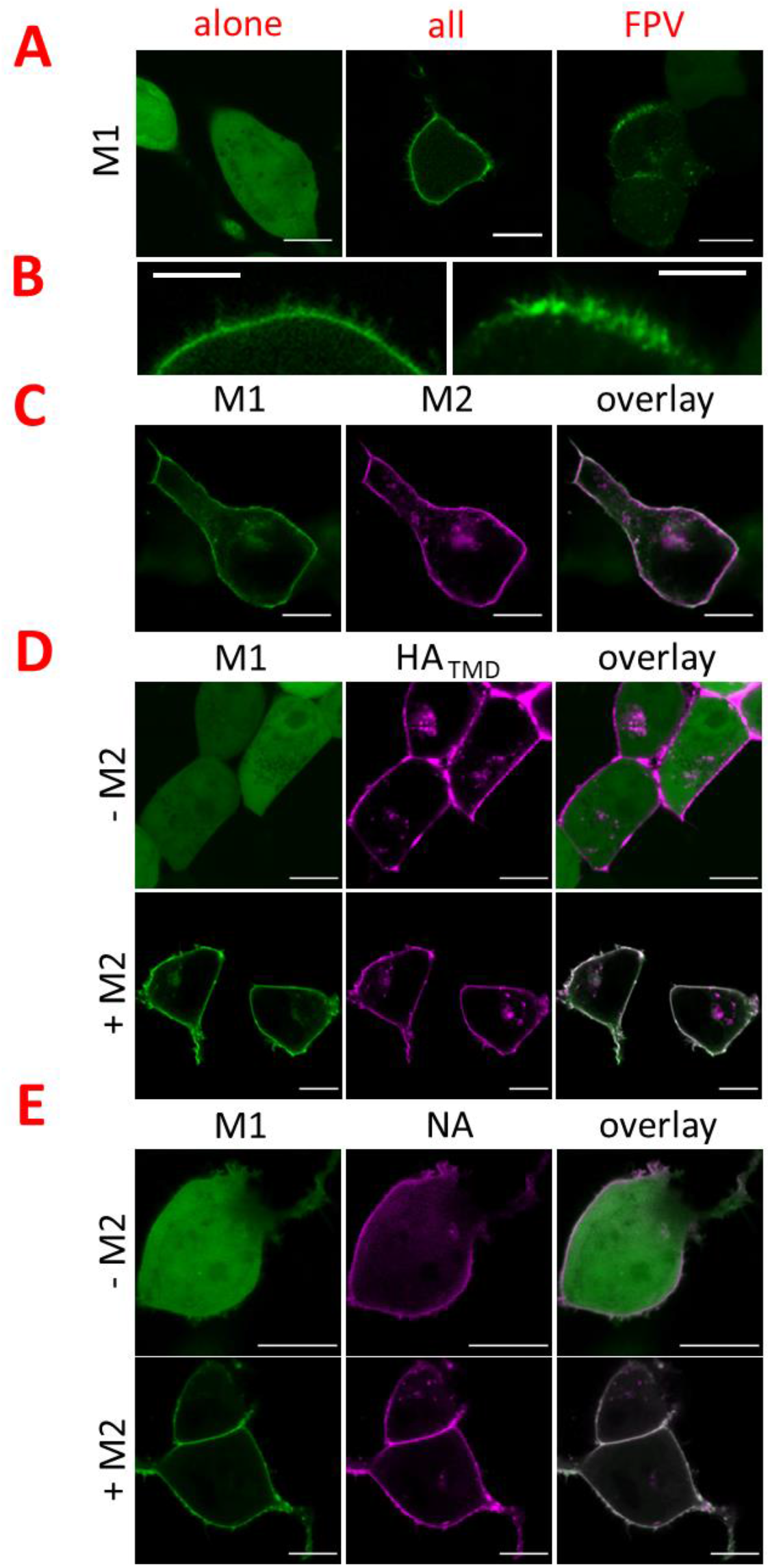
Membrane recruitment of IAV matrix protein 1 (M1) in co-transfected and infected cells. A-B: Representative confocal fluorescence images of HEK293T cells expressing M1-mEGFP (green) from the influenza A/FPV/Rostock/1934 strain (FPV) alone (A, left panel). The same construct was also observed in cells co-transfected with the reverse genetic plasmid system of FPV and unlabeled M2, here called as “all” (A, middle; B, left) and in cells infected with FPV (A, right; B, left). C: Representative confocal fluorescence images of HEK293T cells co-expressing M1-mEGFP (green) and the FPV matrix protein 2 (mCherry2-M2, magenta). The right panels show the two channels merged in a single image. D: Representative confocal fluorescence images of HEK293T cells co-expressing M1-mEGFP (green) and the hemagglutinin (mCherry2-HA_TMD_, magenta) in the absence (upper panels) or in the presence (lower panels) of unlabeled M2. E: Representative confocal fluorescence images of HEK293T cells co-expressing M1-mEGFP (green) and the neuraminidase (NA-mCherry2, magenta) in the absence (upper panels) or in the presence (lower panels) of unlabeled M2. Scale bars represent 10 μm.

Expression of M1-mEGFP alone indicated a homogenous distribution of M1 through the cytosol and the nucleus (Figure 1 A), whereas mEGFP-M1 (i.e., mEGFP fused at the N-terminus) formed large, bright aggregates in the cytosolic region in close proximity to the nucleus (data not shown). The localization of M1-mEGFP was similar to what was previously described for unlabeled M1 (46, 47). Therefore, this construct was used for all further experiments. Upon co-transfection of all other IAV (unlabeled) proteins, a distinct enrichment of M1-mEGFP at the PM was detectable, with the protein being homogeneously distributed (Figure 1 A). A statistical analysis of the frequency of such an occurrence is not trivial since the number of cells effectively transfected with all 9 plasmids is unknown. Nevertheless, a control experiment suggests that most of the successfully transfected cells express several fluorescently tagged proteins at the same time (Figure S2). Also, the probability that a cell expressing M1-mEGFP does not express any other viral protein is estimated to be very low (i.e., <∼1% for a six-plasmid system approximation, Figure S2 B). Therefore, we conclude that the observed enrichment of M1 at the PM is probably due to the presence of at least one other viral protein.

Notably, we observed filamentous structures originating from the PM (Figure 1 B, left, Figure S3 A) that were not present when M1 was substituted by the membrane-anchored mp-mEGFP (Figure S3 A). Cells infected with FPV showed heterogeneous M1 binding to the PM and formation of clusters in almost every cell (i.e. > ca. 90 %) at 24 hpi (Figure 1 A), as previously observed also for unlabeled M1 (15, 16). M1-enriched structures at the PM resembling ruffles were even more evident, compared to the case of the reverse genetic plasmid system (Figure 1 B, right, Figure S3 A). The effectiveness of IAV infection was confirmed via immunofluorescence detection of expressed nucleoprotein (ca. 90 % of infected cells, data not shown).

In order to determine whether M1 localization is determined by the presence of other viral proteins at the PM as previously suggested (15, 16), M1-mEGFP was co-expressed with either mCherry2-M2, mCherry2-HA_TMD_, or NA-mCherry2 (Figure 1 C-E). It is worth noting that these viral proteins are labeled at the extracellular side (so to preserve possible interactions with intracellular M1) and strongly localize at the PM, similarly to their non-fluorescent counterparts (48, 49). Fluorescence microscopy imaging indicated the absence of M1-mEGFP localization at the PM in cells co-expressing this protein with mCherry2-HA_TMD_, NA-mCherry2 constructs (Figure 1D-E) or unlabeled HA or NA (Figure S4 A). On the other hand, upon co-expression of M1-mEGFP with mCherry2-M2, clear colocalization of both proteins at the PM was observed (Figure 1C). Unequivocable association of M1-mEGFP to the PM was observed in circa one quarter of the examined cells and appeared qualitatively correlated with the amount of mCherry2-M2 at the PM (Figure S4 B and C). A quantitative analysis of the correlation between the concentrations of the two proteins at the PM is presented in the following paragraphs.

The membrane distribution of M1-mEGFP was macroscopically homogeneous and no filamentous structures or clustering of M1-mEGFP at the PM were detectable. M2-induced binding of M1-mEGFP to the PM was qualitatively not further influenced by co-expression of mCherry2-HA_TMD_ or NA-mCherry2 (Figure 1 D-E).

In conclusion, M2 seems to be necessary for the recruitment of M1 to the PM. Also, the lateral organization of this protein on the lipid membrane is influenced by the presence of other viral proteins, as observed in infected cells.

### M1 multimeric state at the PM ranges from dimers to large multimers

In order to quantify the concentration-dependent oligomerization of M1, N&B analysis was carried out in infected as well as co-transfected cells (Figure 2). This approach was applied in the past to quantify protein multimerization as a function of local concentration and cellular localization (50, 51). Compared to other methods based on fluorescence fluctuation analysis, N&B provides more representative results in samples characterized by spatial inhomogeneities and slow dynamics (52). The amount of fluorescence signal detected for a single independent protein complex (e.g., a protein dimer) in the unit of time is indicated by the molecular brightness. This parameter is directly connected to the number of fluorophores within such a complex and, therefore, to the multimeric state of the fusion-labeled protein. Specifically, the multimerization can be quantified by normalization of the measured brightness values with the molecular brightness of a monomeric and dimeric reference (see Materials and Methods) (41). To avoid possible interactions between the ectodomain of viral proteins and the solid substrate, we performed all measurements at the equatorial plane of cells rather than the basal membrane (which is often analyzed in the context of N&B studies). Our data show that protein oligomerization can be reproducibly quantified for both PM regions, without substantial differences (Figure S5).

**Figure 2:**
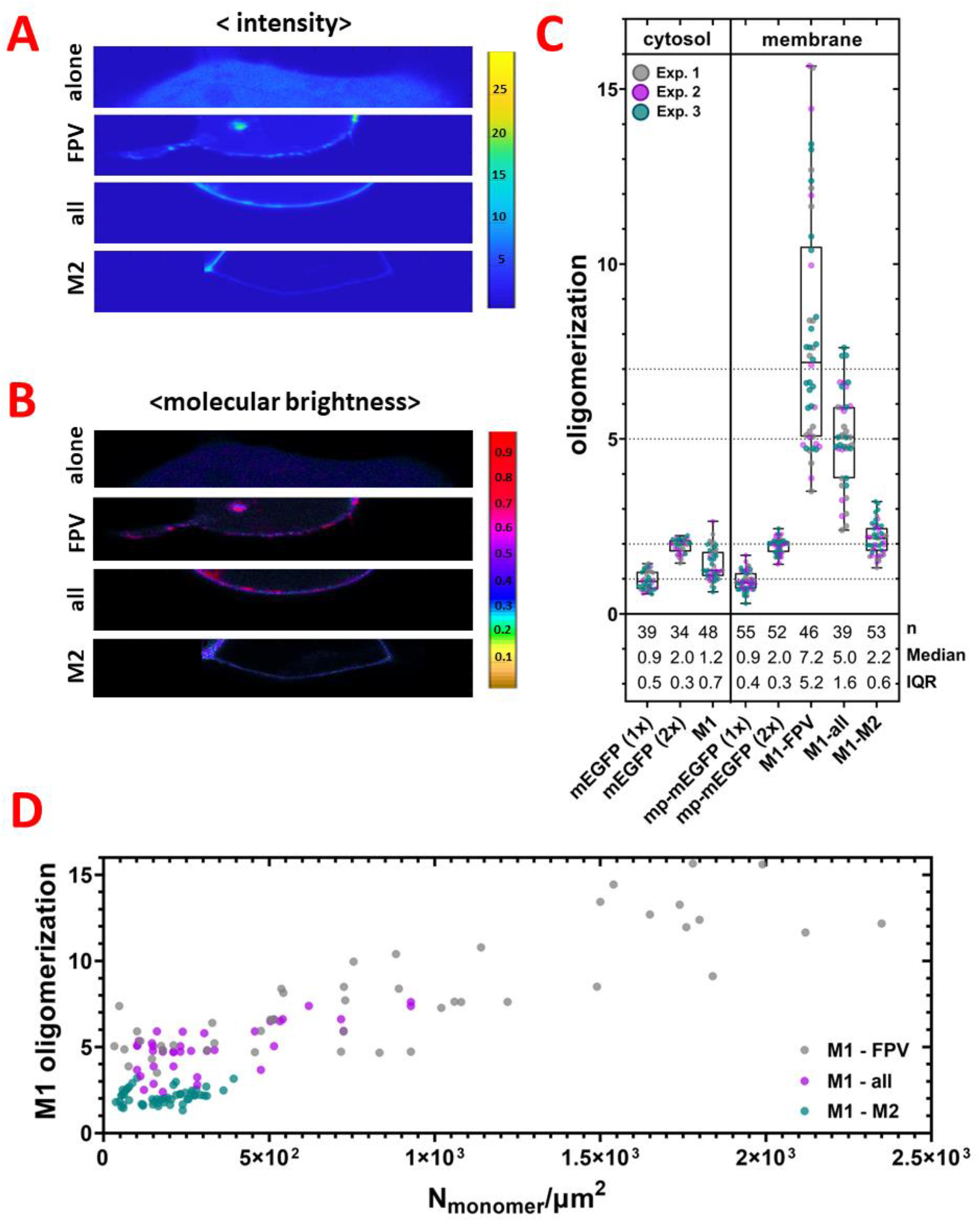
M1 oligomerizes in a concentration-dependent manner. Number and Brightness (N&B) analysis of M1-mEGFP in cells expressing only M1-mEGFP, infected with FPV (“FPV”), co-transfected cells expressing unlabeled M2 and the reverse genetic plasmid system for all other FPV proteins (“all”), or co-transfected cells expressing unlabeled M2 (“M2”). Oligomerization and surface concentration values were obtained as described in the Methods section. A: Representative average intensity maps of M1-mEGFP in HEK293T cells. The average intensity map is visualized via color scale with units photon counts/dwell time. B: Representative brightness-intensity maps corresponding to the images represented in panel A. The images show pixel brightness as pixel color (counts/dwell time per molecule) and mean photon count rate as pixel intensity. C (left): Box plot of single data points from three independent experiments showing the normalized brightness (i.e. oligomerization) for M1-mEGFP and the corresponding controls (i.e., cytosolic monomer mEGFP (1x), cytosolic dimer mEGFP (2x) in the cytosol of HEK293T cells. C (right): Box plot of single data points from three independent experiments showing the oligomerization of M1-mEGFP at the PM of infected (M1-FPV) or co-transfected (M1-all, M1-M2) cells. Oligomerization values for PM-anchored controls (monomer mp-mEGFP(1x), dimer mp-mEGFP(2x)) are also shown. Sample size, median, and interquartile range (IQR) are indicated at the bottom. Horizontal dotted lines corresponding to oligomerization values 1, 2, 5 and 7 are shown as guide to the eye. D: M1-mEGFP oligomerization as a function of surface concentration at the PM (in N_monomer_/μm^2^). The number of measured cells were: M1-FPV (n = 46), M1-all (n = 39), and M1-M2 (n = 53).

The fluorescent construct M1-mEGFP described in the previous paragraph was expressed in HEK293T cells either i) in the presence of unlabeled M2, ii) concurrently with the reverse genetic plasmid system and unlabeled M2, named hereafter “all”, iii) concurrently with FPV infection or iv) alone (Figure 2).

The results shown in Figure 2 A and B indicate that, upon co-expression of M2, M1-mEGFP does not form large complexes, compared to the case in which other viral proteins are present (i.e., in the case of the reverse genetic plasmid system or of infection). In the latter cases, higher intensity and brightness values are in fact observed at the PM. The average intensity and molecular brightness values were calculated at each pixel of ROIs (including e.g., the PM or cytosolic regions) and represented as two dimensional histograms (Figure S6, representative example of data from Figure 2 A and B). The brightness values of M1-mEGFP within each cell were usually symmetrically distributed around their average values for co-transfected cells expressing unlabeled M2, but slightly skewed towards large values in infected cells or cells transfected with the plasmid set “all”. The brightness values of such distributions were then normalized using the corresponding monomer and dimer controls (Figure 2 C). The analysis of cells expressing only M1 indicated that M1-mEGFP in the cytosol has a normalized brightness between 1 and 2 (1.2 ± 0.7, median ± IQR, n = 48 cells). For comparison, the oligomerization state of cytosolic control monomers (mEGFP) and dimers (mEGFP-mEGFP) is also shown. It is worth noting that N&B analysis provides an average oligomerization value in the case of mixtures of different multimeric species (51). Therefore, the measured normalized brightness for cytosolic M1-mEGFP suggests that the protein is present as a mixture of e.g. monomers (ca. 80 %, assuming *p*_*f*_ = 0.7) and dimers (ca. 20 %) at the observed concentrations. M1-mEGFP oligomerization slightly increased upon binding to the PM in the presence of unlabeled M2 (2.2 ± 0.6, median ± IQR, n = 53 cells). M1-mEGFP oligomeric state increased significantly upon co-transfection with all other viral proteins (“all”, 5.0 ± 1.6, median ± IQR, n = 39 cells), or upon infection (7.2 ± 5.2, median ± IQR, n = 46 cells). For comparison, the oligomeric state of control monomers (mp-mEGFP) and dimers (mp-mEGFP-mEGFP) is also shown. Additionally, M1-mEGFP showed a significant concentration-dependent oligomerization behavior in concurrently infected cells and in transfected cells expressing all other viral proteins (Figure 2 D). On the other hand, the oligomerization of M1-mEGFP in co-transfected cells expressing unlabeled M2 seemed to be independent from concentration and stable around values corresponding, in average, to M1-mEGFP dimers. As also evident from Figure 2 D, higher concentrations of M1-mEGFP at the PM were observed in general in infected cells, as well as in co-transfected cells expressing all other viral proteins.

Of note, it must be considered that in infected cells, M1 concentration and oligomerization are underestimated, due to the co-expression of viral unlabeled M1 which might take part in the formation of complexes with M1-mEGFP. Since N&B analysis accounts only for labeled proteins, complexes containing both labeled and unlabeled species will effectively appear as smaller oligomers. Additionally, it is also possible that a precise determination of the multimeric state might be hindered by high protein concentrations at the PM, especially for very large multimers.

In summary, M1-mEGFP forms up to dimers in the cytoplasm or at the PM, upon co-expression of M2. The oligomerization of membrane-bound M1-mEGFP increases dramatically as a function of local concentration in infected cells and, to a minor extent, in cells expressing all other viral proteins via a reverse genetic plasmid system.

### HA and NA do not induce M1 oligomerization

The interaction of M1 with other viral membrane proteins (HA, NA, and M2) is controversially discussed in previous studies (10, 11, 22-24, 30, 31, 35, 36).

To clarify this issue, we performed 2-color sFCCS analysis in HEK293T cells expressing M1-mEGFP in combination with i) mCherry2-M2, ii) mCherry2-HA_TMD_ and unlabeled M2, or iii) NA-mCherry2 and unlabeled M2. As shown for example in Figure 3 A for the case of co-transfected cells expressing M1-mEGFP, mCherry2-HA_TMD_, and unlabeled M2, M1 partitions strongly at the PM in all cases. For sFCCS measurements, the confocal detection volume is scanned in a linear fashion perpendicularly to the PM, as illustrated by the yellow arrow. Following the calculation of ACFs and CCFs, (Figure S7), this approach allows the quantification of the interactions between two differently labeled proteins by calculating the relative cross-correlation (rel. cc), i.e. a measure of the relative abundance of molecular hetero-complexes. Furthermore, from the analysis of the ACF, sFCCS provides quantitative information about diffusion dynamics and, similar to N&B analysis, the average oligomerization state of the monitored proteins.

**Figure 3:**
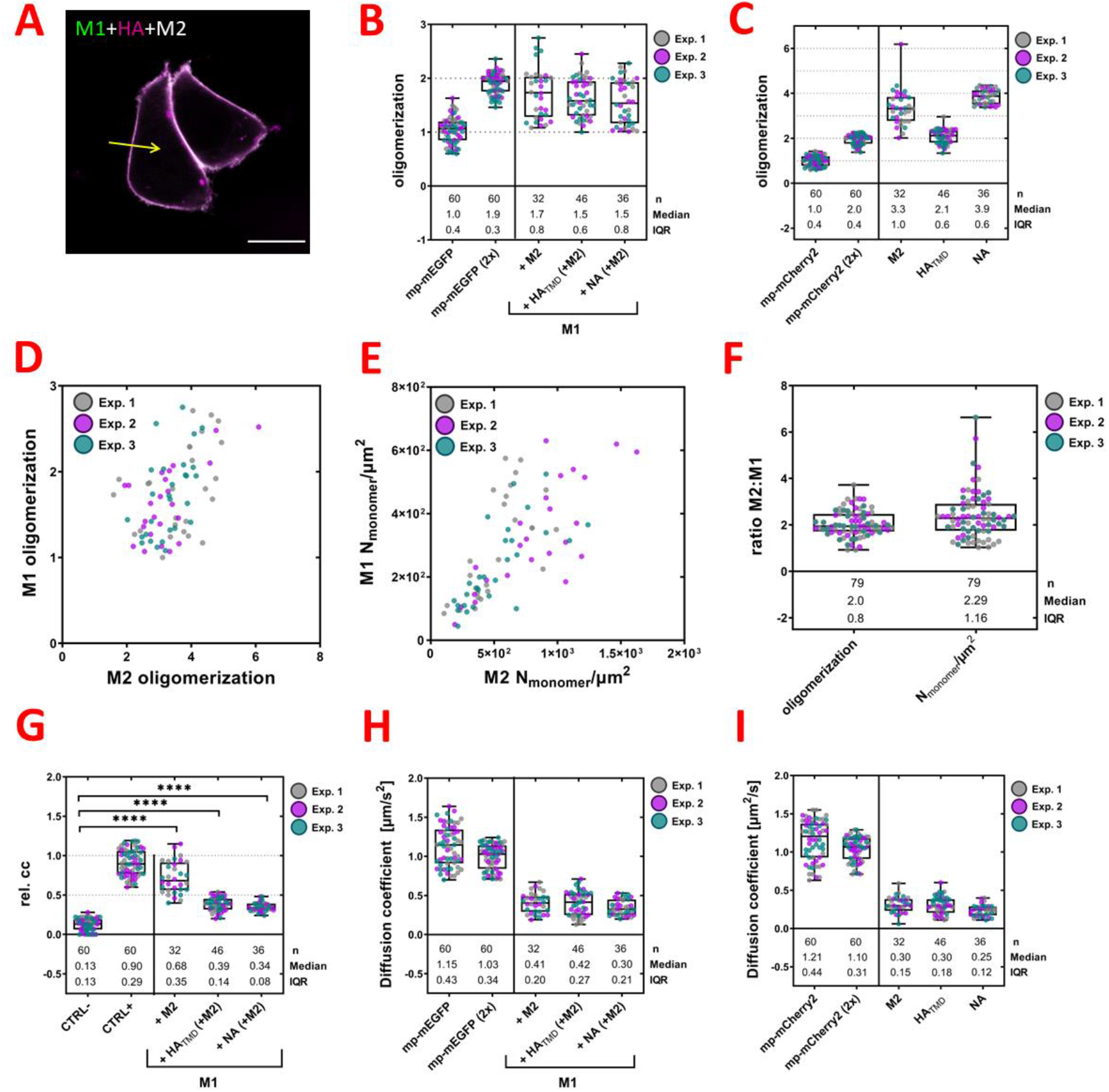
M2 interacts with M1 in a concentration-dependent manner. Scanning fluorescence cross-correlation spectroscopy (sFCCS) of M1-mEGFP in HEK293T cells co-expressing mCherry2-M2, mCherry2-HA_TMD_/M2-untagged, and NA-mCherry2/M2-untagged. Oligomerization, surface concentration (N_monomer_/μm^2^), cross-correlation, and diffusion coefficient (μm^2^/s) values were obtained as described in the Methods section. A: Representative confocal fluorescence image of HEK293T cells co-expressing M1-mEGFP (green), mCherry2-HA_TMD_ (magenta), M2-untagged. Yellow arrow indicates the scanning path used for sFCCS. Scale bar is 10 μm. B: Box plot with single data points from three independent experiments shows the oligomerization of the controls (monomer mp-mEGFP(1x) and dimer mp-mEGFP(2x)), and M1-mEGFP co-expressed with mCherry2-M2, mCherry2-HA_TMD_ /M2-untagged, and NA-mCherry2/M2-untagged. Sample size, median, and IQR are indicated in the graph. C: Box plot with single data points from three independent experiments shows the oligomerization of the controls (monomer mp-mCherry2(1x) and dimer mp-mCherry2(2x)), and the viral surface proteins mCherry2-M2, mCherry2-HA_TMD_, and NA-mCherry2 for the same samples described for panel B. Sample size, median, and IQR are indicated in the graph. D-E: Scatter plots show the oligomerization of M1-mEGFP as a function of the oligomerization of mCherry2-M2 (D), and the surface concentration of M1-mEGFP as a function of the surface concentration of mCherry2-M2 (E). F: Box plot with single data points from three independent experiments shows the ratio of the oligomerization, and the surface concentration of M2:M1. Sample size, median, and IQR are indicated in the graph. G: Box plot with single data points from three independent experiments shows the relative cross-correlation (rel. cc) for the controls (negative control mp-mEGFP(1x)/mp-Cherry2 and positive control mp-mCherry2-mEGFP) and between M1-mEGFP and mCherry2-M2, mCherry2-HA_TMD_, or NA-mCherry2. Cells expressing mCherry2-HA_TMD_ and NA-mCherry2 also expressed unlabeled M2. Sample size, median, and IQR are indicated in the graph. Statistical significance was determined using one-way ANOVA multiple comparison test (**** indicates P < 0.0001 compared to the negative control (CTRL-)). H: Box plot with single data points from three independent experiments shows the diffusion coefficient of the controls (monomer mp-mEGFP(1x) and dimer: mp-mEGFP(2x)), and M1-mEGFP co-expressed with mCherry2-M2, mCherry2-HA_TMD_ /M2-untagged, and NA-mCherry2/M2-untagged. Sample size, median, and IQR are indicated in the graph. I: Box plot with single data points from three independent experiments shows the diffusion coefficient of the controls (monomer mp-mCherry2(1x) and dimer mp-mCherry2(2x)), and the viral surface proteins mCherry2-M2, mCherry2-HA_TMD_, and NA-mCherry2 for the same samples described for panel H. Sample size, median, and IQR are indicated in the graph.

Our results suggest that M1 forms monomers and dimers at the PM, upon co-expression of M2 (1.7 ± 0.8, median ± IQR, n = 32 cells), confirming the results of the N&B experiments (Figure 3 B). For comparison, the oligomerization state of control monomers (mp-mEGFP) and dimers (mp-mEGFP-mEGFP) is also shown. Further, the oligomerization of M1 is not significantly altered by additionally co-expressing the IAV glycoproteins, mCherry2-HA_TMD_ (1.5 ± 0.6, median ± IQR, n = 46 cells) or NA-mCherry2 (1.5 ± 0.8, median ± IQR, n = 36 cells). To verify whether the FP fused to viral glycoproteins alters their quaternary structure, the molecular brightness of mCherry2-HA_TMD_ and NA-mCherry2 was also analyzed and compared to the corresponding controls (Figure 3 C). The HA transmembrane domain construct mCherry2-HA_TMD_ formed in average dimers (2.1 ± 0.6, median ± IQR, n = 46 cells), and NA– mCherry2 formed in average tetramers (3.9 ± 0.6, median ± IQR, n = 36 cells). Both oligomeric states are consistent with those obtained in earlier studies (53, 54). The average oligomerization state of mCherry2-M2 (3.3 ± 1.0, median ± IQR, n = 32 cells) indicated that M2 might be present as a mixture of e.g. dimers and tetramers on the PM, which is consistent with previous results (55). Surprisingly, for all the examined IAV proteins, we observed that their average oligomerization state was not strongly influenced by their local concentration (Figure S8).

It is worth noting that the mCherry2-M2 construct (i.e., with mCherry2 fused to the N-terminus of M2) was newly designed to monitor M1:M2 interactions while avoiding steric hindrance at the cytosolic side of M2. In order to determine whether this fluorescent M2 construct behaves as expected (especially in the context of M1-M2 interactions), we used an alternative strategy to simultaneously express untagged M2 and a membrane marker (mp-mCherry2) via a bi-directional vector system (indicated as M2 ↔ mp-mCherry2) (56). The measured concentration of mp-mCherry2 can be used to estimate the amount of unlabeled M2 in the PM (see Materials and Methods in SI, Figure S9). The correct expression of M2 at the PM was validated by immunofluorescence (Figure S9 C). No significant difference in the oligomeric state of M1-mEGFP as a function of the surface concentration of M2 between both plasmid constructs (i.e., bidirectional M2 and mCherry2-M2) was observed (Figure S9 D). Therefore, only the mCherry2-M2 construct was used for further investigations of M1-M2 interaction.

Notably, the oligomerization of M1-mEGFP was consistently independent from the concentration of mCherry2-M2 at the PM (Figure S9 D) but correlated with the oligomerization state of mCherry2-M2 (Figure 3 D). Also, the concentration of M1-mEGFP at the PM increased with increasing mCherry2-M2 concentration (Figure 3 E). As shown in Figure 3 F, we could finally estimate that both M1-mEGFP concentration at the PM and oligomerization are circa half of what is observed for mCherry2-M2 (M2:M1_oligo.state_: 2.0 ± 0.8, and M2:M1_surface conc._: 2.29 ± 1.16, median ± IQR).

In summary, our results suggest that M1 binds to the PM as dimer upon co-expression of M2. M1-M1 and M1-lipid interactions did not appear to be modulated by the presence of HA or NA.

### M1 strongly interacts with M2 but only weakly associates to HA or NA

Direct information regarding the formation of protein hetero-complexes at the PM can be derived by the analysis of ACFs and CCFs obtained via sFCCS (see previous paragraph). We therefore calculated the rel. cc as a measure of the hetero-interactions between M1-mEGFP and either mCherry2-M2, mCherry2-HA_TMD_, or NA-mCherry2 (Figure 3 G, and S3). Two interacting molecules diffusing together through the observation volume as a complex will give rise to a positive rel. cc that can be quantified by the amplitude of the cross-correlation curve (Figure S7 B). Low rel. cc indicates the absence of interaction between the observed proteins (see e.g., Figure S7 A). However, due to incomplete maturation of the fluorescent proteins and the partial overlap of the confocal volumes in both channels, the maximum achievable rel. cc value is lower than 1. For example, a tandem of mp-mCherry2-mEGFP used here as a positive control for rel. cc displayed a rel. cc of 0.90 ± 0.29 (median ± IQR, n = 60 cells). As expected, we detected a very low rel. cc (0.13 ± 0.13, median ± IQR, n = 60 cells) in negative control experiments (i.e., in samples of co-transfected cells expressing mp-mEGFP and mp-mCherry2). As shown in Figure 3 G, a rel. cc of 0.7 ± 0.4 (median ± IQR, n = 32 cells) was measured for M1-mEGFP and mCherry2-M2. This value is significantly higher than the negative control and close (ca. 80 %) to that obtained for the positive control, suggesting very strong association of M1-mEGFP with mCherry2-M2. Assuming a very simple scenario consisting e.g. of M1 dimers, M2 tetramers and 2:4 M1-M2 complexes, all detectable with p_f_=1, ca. 80% of M1 molecules appear to be in complex with M2.

On the other hand, the obtained rel. cc values for M1-mEGFP with either mCherry2-HA_TMD_, or NA-mCherry2 (rel. cc(M1,HA_TMD_) = 0.39 ± 0.14, n = 46 cells; rel. cc(M1,NA) = 0.34 ± 0.08, n = 36 cells, median ± IQR) were lower but still significantly higher than the negative control. It is worth noting that such measurements could only be performed in the presence of unlabeled M2 since, without this third protein, no localization of M1-mEGFP at the PM could be observed (see previous paragraphs). The observed rel. cc values indicate a weak interaction between M1-mEGFP and the glycoproteins mCherry2-HA_TMD_, and NA-mCherry2. In the simple approximation of p_f_=1 and constant multimerization, independently from the participation in complexes, ca. 40% of M1 molecules appear to be associated with the viral glycoproteins. To further investigate this issue, we quantified the interaction between M1 and the glycoproteins also in infected cells. To this aim, we performed ccN&B in cells infected with FPV and, additionally, co-transfected with M1-mEGFP and either mCherry2-HA_TMD_ or NA-mCherry2 plasmids. Similar to sFCCS, ccN&B can be used to quantify the rel. cc between different FPs, especially in samples characterized by slow dynamics (52). Scanning FCCS measurements of M1-mEGFP in infected cells did not provide in fact reproducible results (data not shown). As shown in Figure S10, the rel. cc values determined by ccN&B in infected cells for M1-mEGFP and mCherry2-HA_TMD_ (rel. cc(M1,HA_TMD_) = 0.31 ± 0.10, n = 21 cells, median ± IQR), as well as for M1-mEGFP and NA-mCherry2 (rel. cc(M1,NA) = 0.28 ± 0.08, n = 22 cells, median ± IQR) were roughly comparable to the rel. cc values obtained in non-infected cells, as measured via sFCCS (Figure 3 G). A more precise quantification is complicated in this case by the presence of non-fluorescence viral proteins and unknown stoichiometry of the investigated molecular complexes.

Finally, we quantified protein dynamics by fitting a two-dimensional diffusion model to the ACF data (Figure 3 H and I, S7). Knowing the size of the observation volume, it is possible to obtain diffusion coefficients of the proteins (D in μm/s^2^, see Materials and Methods). Protein diffusion depends in general on the size of the protein complex and on protein-membrane interactions. The diffusion coefficients measured for M1-mEGFP at the PM (D = 0.3 - 0.4 μm/s^2^, Figure 3 H) were lower than those of the monomer control (D = 1.1 ± 0.4 μm/s^2^, median ± IQR, n = 60), and similar to the diffusion coefficient of the IAV integral surface proteins mCherry2-M2, mCherry2-HA_TMD_, and NA-mCherry2 (indicated in Figure 3 I).

Taken together, our data indicate that M1 strongly interacts with M2. On the other hand, a relatively small amount of complexes containing M1 together with HA or NA was detected.

### Non-specific M1 recruitment to the PM is sufficient for the establishment of M1-HA interaction

To investigate the origin of the interaction between M1 and HA (or NA) which was observed in cells additionally expressing M2, we artificially induced M1 binding to the PM. These experiments were performed to test the hypothesis that M1 is recruited (by M2) to the PM, where it can then interact with other membrane proteins (independently from the specific protein that first induced M1-PM binding).

Specifically, we designed two M1 constructs in which the protein was modified by myristoylation and palmitoylation (mp-M1-mEGFP) and, additionally, with a poly-lysine motif (mp-KrΦ-M1-mEGFP), as shown in Figure 4 A. The underlying idea is that the additional targeting sequences direct M1 specifically to lipid ordered “raft” domains (myristoyl-palmitoyl anchor (57)) or to regions containing acidic lipids (poly-lysine motif) in the PM, as supported by previous studies (27, 58-60). M1 localization within specialized PM domains might be indeed relevant, since the viral envelope proteins HA and NA were previously reported to localize in lipid “rafts”, whereas M2 was observed at the edges between ordered and disordered domains (7, 14).

**Figure 4:**
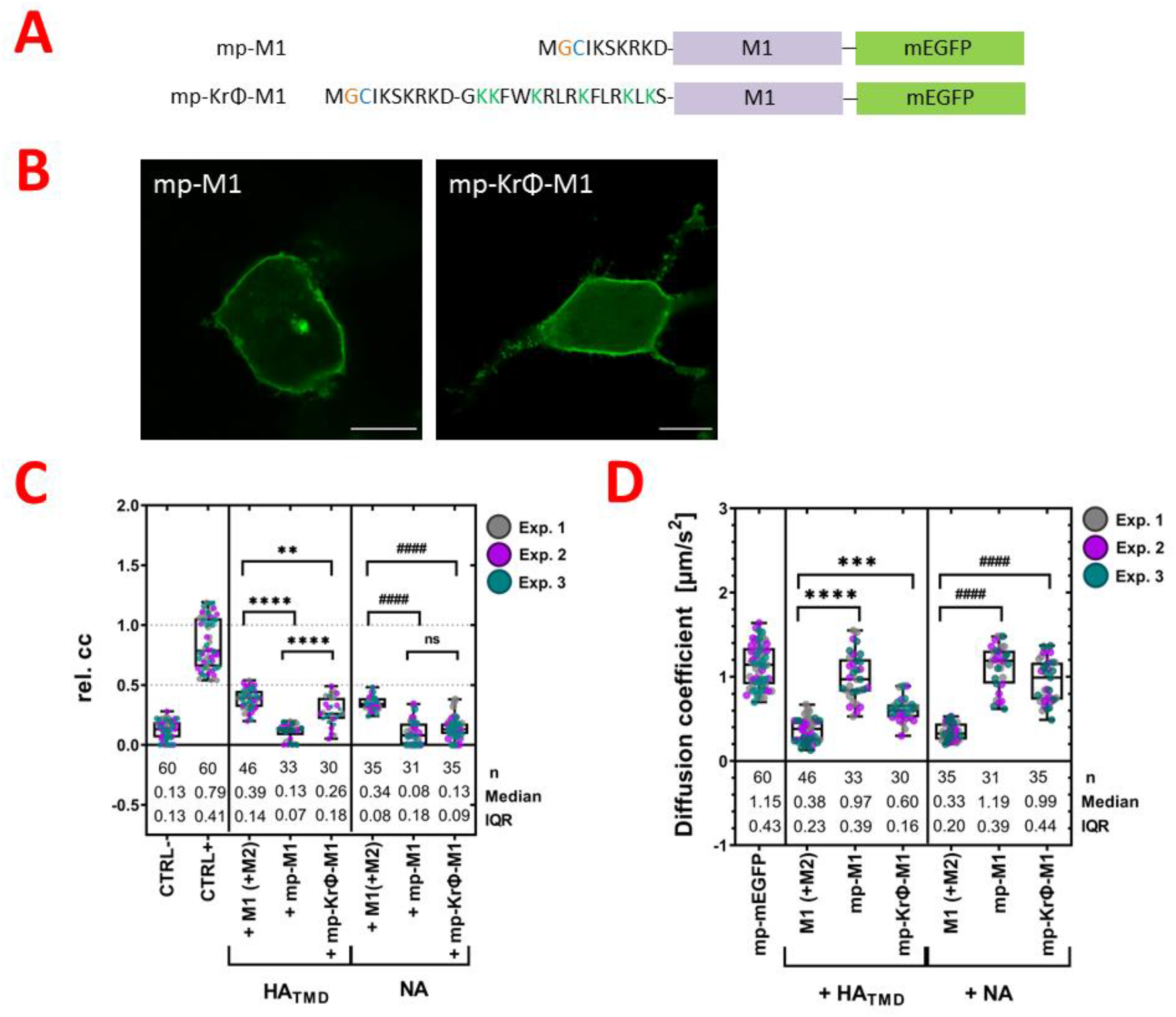
HA interacts with a membrane-associated M1 construct. A: Schematic diagram of M1 constructs with N-terminal PM-targeting sequences. One construct has a myristoylation (orange) and palmitoylation (blue) motif (mp-M1-mEGFP), and the other on has an additional poly-lysine motif (green letters, mp-KrΦ-M1-mEGFP). B: Representative M1 subcellular localization images in transfected HEK293T cells expressing mp-M1-mEGFP (left side), or mp-KrΦ-M1-mEGFP (right side). Scale bars represent 10 μm. C: Box plot with single data points from three independent experiments shows the cross-correlation for the controls (negative control mp-mEGFP(1x)/mp-mCherry2(1x) and positive control mp-mCherry2-mEGFP), and between M1-mEGFP (or mp-M1-mEGFP, or mp-KrΦ-M1-mEGFP) and mCherry2-HA_TMD_, or NA-mCherry2. Sample size, median, and IQR are indicated in the graph. Statistical significance was determined using one-way ANOVA multiple comparison test (** indicates p < 0.01, **** indicates p < 0.0001 compared to M1-mEGFP/ mCherry2-HA_TMD_; ^####^ indicates p < 0.0001 compared to M1-mEGFP/ NA-mCherry2, ns indicates not significant). D: Box plot with single data points from three independent experiments shows the diffusion coefficient of the monomer control (mp-mEGFP), and M1-mEGFP, mp-M1-mEGFP, and mp-KrΦ-M1-mEGFP co-expressed with mCherry2-HA_TMD_, or NA-mCherry2. Sample size, median, and IQR are indicated in the graph. Statistical significance was determined using one-way ANOVA multiple comparison test (*** indicates p < 0.001, **** indicates p < 0.0001 compared to M1-mEGFP/ mCherry2-HA_TMD_; ^####^ indicates p < 0.0001 compared to M1-mEGFP/ NA-mCherry2).

First, we verified the sub-localization of the two new constructs in transfected HEK293T cells. Both, mp-M1-mEGFP and mp-KrΦ-M1-mEGFP, were efficiently trafficked to the PM (Figure 4 B). Next, we examined the rel. cc between these two constructs and mCherry2-HA_TMD_, as well as NA-mCherry2 (Figure 4 C) in co-transfected cells. The obtained rel. cc values (indicated in Figure 4 C) for mp-M1-mEGFP with mCherry2-HA_TMD_ or NA-mCherry2, as well as mp-KrΦ-M1-mEGFP with NA-mCherry2, were similar to those of the negative rel. cc control. These results indicate that NA-mCherry2 does not significantly interact with any of the modified membrane-associated M1 constructs. Also, mCherry2-HA_TMD_ does not seem to interact with the supposedly lipid raft-associated mp-M1-mEGFP. In contrast, a reproducible interaction between mp-KrΦ-M1-mEGFP and mCherry2-HA_TMD_ (rel. cc(mp-KrΦ-M1,HA_TMD_) = 0.26 ± 0.18, n = 30 cells, median ± IQR) was observed. Notably, the rel. cc value observed in this case was significantly lower than the one obtained in the context of the interaction between (wildtype) M1-mEGFP and mCherry2-HA_TMD_, in the presence of M2. Next, we calculated the surface concentration of each protein and plotted the cross-correlation values against the surface concentration, as well as the ratio of the concentration between the protein pairs (Figure S11). This analysis was performed to exclude that the obtained rel. cc values are influenced by the surface concentration of the proteins or the expression ratio between the proteins. No concentration-dependency of the rel. cc for all pairs was observed.

Finally, we quantified the diffusion dynamics of the examined protein constructs (Figure 4 D). The obtained diffusion coefficient values (shown in Figure 4 D) for mp-M1-mEGFP in the presence of mCherry2-HA_TMD_ or NA-mCherry2 were similar to those of the monomer control (mp-mEGFP). A similar observation was made for mp-KrΦ-M1-mEGFP in the presence of NA-mCherry2. The fact that these M1 constructs diffuse as fast as a lipid-anchored protein (rather than a membrane-spanning protein, see Figure 3 I) suggests the absence of significant interactions/co-diffusion of M1 with mCherry2-HA_TMD_ or NA-mCherry2. For comparison, the diffusion coefficients of M1-mEGFP in the presence of M2 and one glycoprotein are also reported in Figure 4 D (D = 0.38 ± 0.23 μm/s^2^, median ± IQR, n = 46, when co-expressed e.g. with mCherry2-HA_TMD_). This result is comparable to the diffusion coefficient of mCherry2-M2 (D = 0.30 ± 0.15 μm/s^2^, median ± IQR, n = 46, Figure 3 I). Interestingly, the diffusion coefficient for mp-KrΦ-M1-mEGFP (D = 0.60 ± 0.16 μm/s^2^, median ± IQR, n = 32) co-expressed with mCherry2-HA_TMD_ was slightly lower than that measured for the monomer control, although still higher than the one measured for M1-mEGFP in the presence of unlabeled M2. It is also worth noting that the distribution of diffusion coefficient values for the above-mentioned sample appears to slightly deviate from a normal distribution (Kolmogorov-Smirnov test P value= 0.0356). The reason for this deviation is not clear at this point but one possible cause might be, for example, the occasional presence of cytosolic signal (see e.g. Figure 4 B) weakly interfering with measurements at the PM.

In conclusion, NA-mCherry2 does not exhibit significant cross-correlation or co-diffusion with neither of the “artificially” PM-associated M1 proteins. In contrast, mCherry2-HA_TMD_ appears to interact with M1 depending on the specific way in which the latter is anchored to the PM.

### A potential M2 binding site is located in the N-domain (aa 1-67) of M1

An interaction site for M2 has not been yet identified within M1. Therefore, we created different M1 constructs for the expression of specific protein subdomains, in order to locate a potential M2 binding site (Figure 5 A). The truncated M1 constructs encoded i) the N- and M M-domains (NM1, amino acids 1–164), ii) the N-terminus domain including the linker region (NM1, amino acids 1–86), iii) only the N-domain (NM1, amino acids 1–67) or iv) the M1 C-domain (CM1, amino acids 165–252). A mEGFP was fused to the C-terminal site of each M1 variants. Moreover, a well conserved amino acid sequence in the cytoplasmic C-terminal tail of M2 at the position 71 and 73 was previously shown as an interaction site for M1 (10). Hence, we generated a substitution mutant of M2 (M2_mut_) in which the triplet sequence (71 – SMR – 73) was replaced by alanine (Figure 5 A).

**Figure 5:**
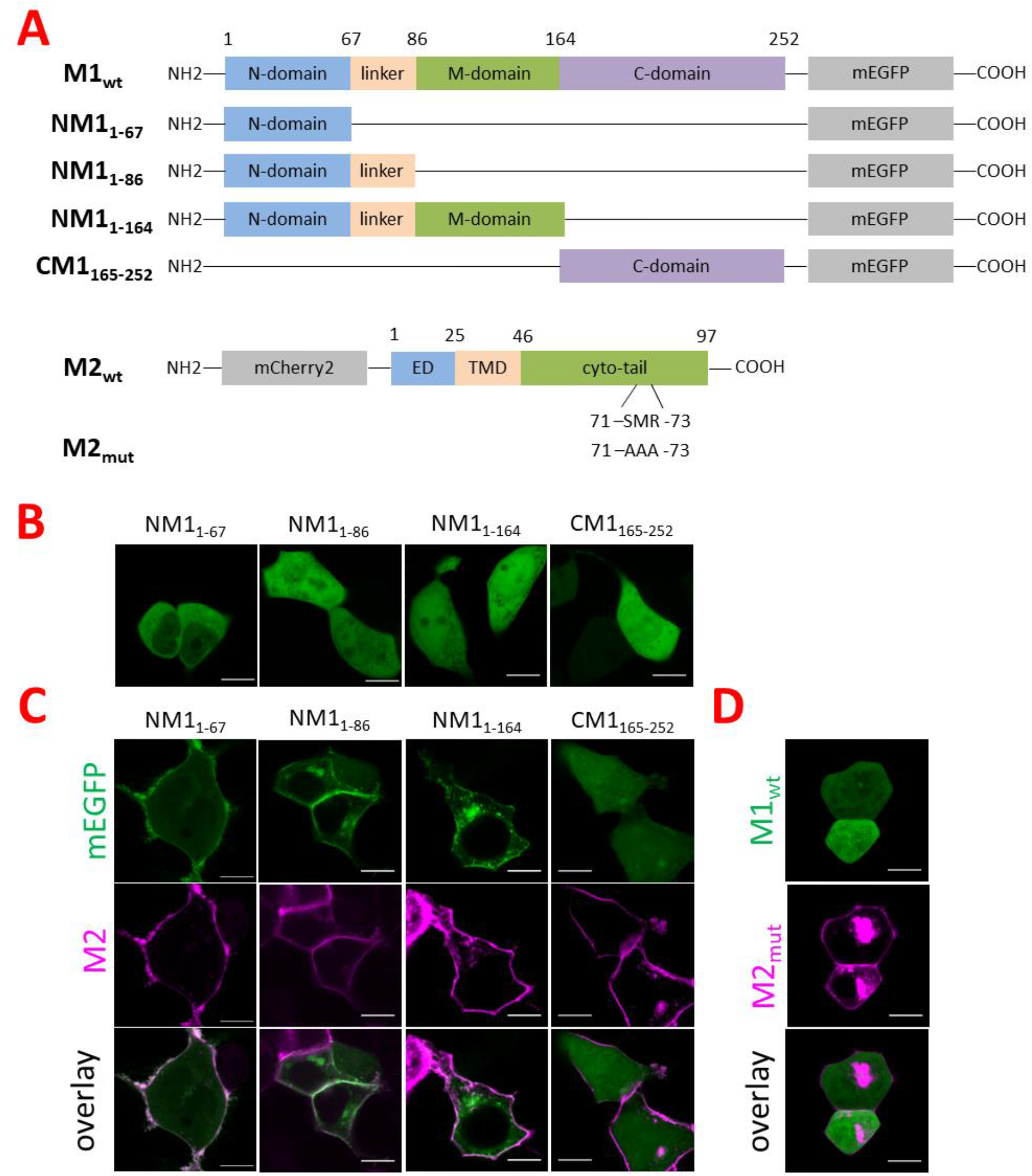
M2 binding site on M1 is located in the N-terminus domain. A: Schematic diagram of different M1 and M2 expression constructs. On top, M1 constructs showing the wildtype and the truncated M1 variants with their domains: N-terminus domain (N-domain, aa 1-67, blue), linker region (orange, aa 68-86), middle domain (M-domain, aa 87-164, green), and C-terminus domain (C-domain, aa 165-252, purple). A mEGFP was fused to the C-terminus of each protein construct. On the bottom, M2 constructs showing the wildtype and the M2 mutant (71-SMR-73 was replaced by three alanine) with their domains: ectodomain (ED, aa 1-25, blue), transmembrane domain (TMD, aa 26-46, orange), cytoplasmic tail (cyto-tail, aa 47-97, green). Each construct had a mCherry2 fused to the N-terminal site of M2. B: Representative confocal fluorescence images of HEK293T cells expressing truncated M1-mEGFP variants: NM1_1-67_, NM1_1-86_, NM1_1-164_, and CM1_165-252_. C: Representative confocal fluorescence images of HEK293T cells expressing truncated M1-mEGFP variants: NM1_1-67_, NM1_1-86_, NM1_1-164_, and CM1_165-252_ (green) in the presence of wildtype mCherry2-M2 (magenta). (D) Representative confocal fluorescence images of HEK293T cells expressing wildtype M1-mEGFP (green) with mCherry2-M2 mutant (M2_mut_, magenta). Scale bars represent 10 μm.

To verify whether the truncated M1-mEGFP constructs are altered in their subcellular localization, we observed them in HEK293T in the absence of mCherry2-M2. All truncated M1-mEGFP variants showed a similar subcellular localization to the wildtype M1-mEGFP (Figure 1 A, and 5 B). Next, all truncated M1-mEGFP constructs were co-expressed with mCherry2-M2 in HEK293T cells. All N-terminus M1 variants were recruited to the PM whereas the C-terminus M1 construct showed still a homogeneous distribution in the cytoplasm (Figure 5 C). The percentages of cells with M1 at the PM for the NM1 variants were similar as observed for the M1 wildtype in co-expression with mCherry2-M2 (Figure S4 C). These results indicated that the M2 binding site might be pinpointed to the N-terminal domain of M1 and, specifically, to the amino acids 1-67. Furthermore, no recruitment of M1 wildtype was observed upon a co-expression with the mCherry2-M2_mut_ (Figure 5 D). Based on this result, we could confirm that the recruitment of M1 to PM occurred via a specific interaction of M1 with the amino acid sequence (71 – SMR – 73) on M2.

## Discussion

The role of M1 in IAV assembly is of fundamental importance, as it is now understood that this protein connects together the viral envelope, its membrane proteins (HA, NA, and M2) and the genome (61). It has been suggested that interactions of M1 with the viral glycoproteins (e.g. HA) drive M1 localization to the PM of infected cells (10, 11, 22-24), but other studies reported conflicting data regarding the interaction of M1 with HA and NA (30, 31, 35, 36). Such findings are mostly based on biochemistry approaches providing indirect interaction data (7, 62).

Therefore, in order to quantify protein-protein interactions directly at the PM of living cells, we performed fluorescence fluctuation spectroscopy experiments under physiological conditions. Such experimental approaches (i.e., sF(C)CS and (cc)N&B) provide information regarding the oligomeric state, surface concentration, hetero-interactions and dynamics in complex biological systems (51, 52, 63-66).

To this aim, we selected HEK293T cells as a cellular model because they are often used for reverse genetic virus production (39, 40, 67) and were shown to be appropriate for IAV protein expression (17, 41, 42). Additionally, we produced and tested several fluorescent IAV protein constructs. Of note, the fluorescent NA construct designed in this study allowed for the first time the investigation of the interaction between this IAV glycoprotein and other viral proteins directly in living cells. It is worth noting that incorporating fluorescent fusion tags might have an impact in general on the localization, function, and conformation of the protein of interest (68, 69). For example, our control experiments showed that the cellular distribution of M1 with an mEGFP fused to its C-terminus was similar to that of unlabeled M1 (46, 47), whereas an N-terminally fused mEGFP M1 variant seemed to have transport failures which are probably caused by steric hindrance between fluorophore and signal peptide (47). On the other hand, the fluorescent constructs used to investigate the viral envelope proteins (HA, NA, and M2) were all localized at the PM, similar to the corresponding non-fluorescent proteins (48, 49), and yielded the expected oligomerization state (41, 42, 53, 54). For example, our results are compatible with the presence of NA tetramers and mixtures of M2 dimers and tetramers (Figure 3 C), in agreement with previous data (55, 70). Furthermore, we also demonstrated that the protein-protein interactions investigated here (e.g. between M1-mEGFP and mCherry2-M2) are specific and analogous to those observed in other contexts (10), as shown by mutagenesis experiments (Figure 5 D) and using unlabeled interaction partners (Figure S1).

In order to identify the minimum requirement for M1 association to the PM, we observed cells expressing different combinations of viral proteins. First, we confirmed that M1 does not bind to the PM in the absence of other viral proteins, despite the observation of strong lipid-protein interactions previously observed in model membrane systems (17-20). Surprisingly, we did not observe any recruitment of M1 to the PM in the presence of HA or NA (Figure 1). It is worth noting that several studies proposed that M1 interacts with the cytoplasmic tails of HA or NA (23, 30, 71-74), but our direct observations in living cells strongly suggest that the two IAV glycoproteins are not able to recruit M1 to the PM by themselves. It is unlikely that the lack of interaction might be a simple consequence of the presence of fluorescent labels, since HA and NA are labeled at the extracellular side. Also, the same M1-mEGFP strongly associates with the PM in the presence of M2. This result is in agreement with previous studies indicating that M1-M2 interactions affect M1 localization and drive virus assembly (10, 11, 27, 75, 76). For the first time, we could provide direct experimental evidence that the M2-binding region is located within the first 67 amino acids of M1 (Figure 5). Also, thanks to the application of quantitative fluorescence microscopy methods, we could additionally prove that M1 and M2 do not simply colocalize at the PM but rather form complexes. This conclusion is supported by the similar diffusion dynamics observed for M1 and M2 (i.e. diffusion coefficients typical of trans-membrane proteins rather than membrane-associated proteins, Figure 3 H) and by the significant degree of cross-correlation between the signals of the two proteins (Figure 3 G). Due to the lack of strong intracellular co-localization (Figure 1 C and 5 C), we hypothesize that M1 diffuses freely in the cytoplasm and M1-M2 interaction occurs directly at the PM. M1-M2 complexes appear to consist, in average, of 1-2 M1 and 2-4 M2 molecules (Figure 3). Assuming that each M2 monomer has a binding site for M1, the observed 1:2 stoichiometry suggests that the M1 binding might be limited for example by steric constraints or competition with other binding partners of M2 (e.g., LC3 (42)). Furthermore, in the simple approximation of M1 dimers, M2 tetramers, and 2:4 M1-M2 complexes being associated to the PM, our cross-correlation measurements indicate that ca. 80 % of M1 is indeed complexed to M2. The remaining amount of PM-associated M1 might interact with e.g. acidic lipids at the PM (19, 20) but, of note, we never observed any significant degree of M1 localization at the PM in the absence of M2. This finding puts forward the hypothesis that M2-M1 complex formation might facilitate the interaction of M1 with other membrane components. This mechanism might also explain previous findings indicating the presence of HA and M1 in the same membrane fractions (22, 23) or within the same region in the PM (15). Accordingly, we have observed that in the presence of M2 (i.e. M1 being efficiently recruited to the PM) there is a significant (although modest) interaction between M1 and the glycoproteins HA or NA (Figure 3 G). On one hand, it is possible that e.g. M1-HA interactions are not direct but, rather, mediated by M2 (15). Alternatively, it is possible that, while M2 is needed for the initial recruitment of M1 to the PM, M1-M2 interactions are not long-lived and can be partially replaced for example by M1-HA interactions. In this case, M2 might induce interactions between M1 and other membrane components by e.g. increasing M1 local concentration in specific PM regions or stabilizing a certain geometric configuration of M1. Based on control experiments monitoring M1-HA/NA interactions as a function of local protein concentration (Figure S11), a prominent role of concentration seems unlikely though. To evaluate whether M2 is strictly needed for HA-M1 interactions, we performed sFCCS experiments in which M1 was artificially anchored to the PM (Figure 4). In this case, depending on the specific lipid anchor, we were able to observe M1-HA interactions also in the absence of M2, thus indicating that i) the latter protein is not always required as a bridge between M1 and IAV glycoproteins and ii) the lipid environment plays a role in the establishment of interactions among IAV proteins. Of interest, it was shown that HA is associated to specific lipids, such as PI(4,5)P2 (14, 49) and this observation might provide a molecular mechanism explaining our observation of non-negligible M1-HA interactions, in the case that M1 was artificially anchored to the membrane via lipidation and, additionally, a polybasic motif. The degree of association between HA and mp-KrΦ-M1-mEGFP appeared to be between that observed for wt M1 and that observed for mp-M1-mEGFP, as supported by the observation of intermediate cross-correlation values (Figure 4 C) and diffusion dynamics (Figure 4 D).

The observation that one single IAV membrane protein (i.e. M2) is sufficient for the recruitment of M1 to the PM prompted us to investigate whether M1-M2 interaction is also sufficient for the initiation of the large-scale M1 multimerization associated with IAV assembly (17). Our experiments clearly demonstrate that this is not the case, since M1 remains, in average, mostly dimeric when bound to the PM in the presence of M2 (Figure 2 C). On the other hand, in the presence of all the other viral proteins, M1 formed larger multimers (up to 5-10 monomers). This effect does not seem to be a direct consequence of the presence of HA or NA alone (Figure 3 B) and is even stronger in infected cells. It is worth noting that the formation of very large multimers of M1-mEGFP in infected cells might be partially due to i) higher M1 concentrations at the PM or ii) the presence of unlabeled M1 molecules which more efficiently support protein-protein interactions. It was in fact reported that fluorescent viral proteins might not be able to oligomerize on a very large scale (65). Alternatively, other viral proteins or altered lipid metabolism (and, consequently, modification of PM composition) in infected cells might play a role and these possibilities are currently under investigation.

In conclusion, our study sheds light on the very first steps in IAV assembly. According to our results, the main role of M2 in this context is to recruit M1 to specific regions of the PM. This is in agreement with previously proposed models according to which M2 chaperones M1 to the PM (77) and, specifically, to interface regions between “raft” and “non-raft” domains (14, 16) or domains enriched in negatively-charged lipids (17). In further steps, M1 can then interact with lipids and other viral proteins and such interactions might be involved in the formation of larger protein complexes eventually leading to IAV capsid assembly.

## Abbreviations

ACF: autocorrelation function
AF488: Alexa Fluor®488
CCF: cross-correlation function
(cc)N&B: (cross-correlation) number and brightness
FP: fluorescence protein
FPV: influenza A/FPV/Rostock/1934 virus mutant 1
HA: hemagglutinin protein
IAV: influenza A virus
M1: IAV matrix protein 1
M2: IAV matrix protein 2
mEGFP: monomeric enhanced green fluorescent protein
mp: myristoylated and palmitoylated
NA: neuraminidase protein
p_f_: fluorescence probability
PM: plasma membrane
rel. cc.: relative cross-correlation
sFCCS: scanning fluorescence cross-correlation spectroscopy
vRNPs: viral ribonucleoproteins

## Author contributions

Research planning, A.P. and S.C; investigation, A.P.; data analysis, A.P.; writing-original draft preparation, A.P. and S.C.; writing-review and editing, V.D. and S.B.; software, V.D. and S.C.; supervision, S.C.; funding acquisition, S.C.

## Acknowledgments

This work was supported by the German Research Foundation (DFG grant #254850309 to S.C.). The authors thank Andreas Herrmann for reading the manuscript and providing useful feedback.

## Supporting citations

References (78-86) appear in the Supporting Information.

