## Supplementary Info for "Influenza A M2 recruits M1 to the plasma membrane: a fluorescence fluctuation microscopy study"

### Supplemental Information

### Supplemental Materials and Methods

#### Plasmids and cloning.

For the cloning of all following constructs, standard PCRs with custom-designed primers were performed, followed by digestion with FastDigest restriction enzymes and ligation with T4-DNA-Ligase according to the manufacturer's instructions. All enzymes and reagents were purchased from Thermo Fisher Scientific (Waltham, MA, USA) and primers were acquired from Sigma Aldrich trademark of Merck KGaA (Darmstadt, Germany). The correctness of each construct was verified by Sanger sequencing (LGC Genomics GmbH, Berlin, Germany).

The original plasmid encoding FPV matrix protein 1 (M1) with EYFP fused to the N-terminus or C-terminus of M1 (EYFP-M1, M1-EYFP) was a kind gift of Michael Veit (Free University, Berlin, Germany) (1). A monomeric variant of EGFP, containing the A206K mutation (2), was inserted into both M1 constructs by digestion of mEGFP-C1 (gift from Michael Davidson, Addgene plasmid #54759) with AgeI and BsrGI. The modification of M1-mEGFP with an N-terminal mp-signal (amino acids MGCIKSKRKDNLNDDEPV, mp-M1-mEGFP) or an N-terminal mp-signal with an additional polybasic sequence (amino acids MGCIKSKRKDGKKFWKRLRKFLRKLKS, mp-KrΦ-M1-mEGFP) were introduced by PCR with primers encoding the additional amino acids. For the construction of mp-M1-mEGFP and mp-KrΦ-M1-mEGFP, the PCR products were subcloned into mEGFP-N1 (gift from Michael Davidson, Addgene plasmid #54767) with XhoI and EcoRI. The truncated M1 sequences encoding the M1 N-terminus (NM1, amino acids 1–67; 1-86; 1-164) or the M1 C-terminus (CM1, amino acids 165–252) were amplified from the plasmid M1-mEGFP, and subcloned into mEGFP-N1 by using the restriction endonucleases XhoI and EcoRI, yielding plasmids NM1(1-67)-mEGFP, NM1(1-86)-mEGFP, NM1(1-164)-mEGFP, and CM1(165-252)-mEGFP.

An untagged FPV M2 construct was cloned by amplifying M2 sequence from FPV-M2-EYFP (a kind gift from Michael Veit), and cloned into pcDNA3.1+ (Thermo Fisher Scientific, Waltham, MA, USA , #V79020) via restriction with HindIII and EcoRI. The plasmid for bi-directional expression was a gift from Katja Arndt (University of Potsdam, Potsdam, Germany) and contains the two promoters TTC31 and CCDC142, allowing for simultaneous expression of the encoded genes (3). For the calibration of the relative expression level, mp-EGFP and mp-mCherry2 were amplified and cloned into the two expression cassettes flanked by restriction sites BamHI/EcoRI and SacI/KpnI respectively, to obtain mp-mEGFP ↔ mp-mCherry2. A construct with mp-mCherry2 and mp-mEGFP cloned into BamHI/EcoRI and SacI/KpnI restriction sites, respectively, was also produced (mp-mCherry2 ↔ mp-mEGFP). Unlabeled M2 sequence was amplified and separately cloned into the BamHI/EcoRI cassette to produce M2 ↔ mp-mCherry2. Site-directed mutagenesis in order to replace the amino acids of the M1-binding site in M2 (amino acids 71–73, SMR) by alanine residues was performed by two-step overlap-extension PCR of the plasmid mCherry2-M2, yielding the plasmid mCherry2-$M2_{mut}$. The plasmid mCardinal-M2 was cloned based on the previously described mCherry2-M2 (4) by using mCardinal-C1 (gift from Michael Davidson, Addgene plasmid #54590).

The FPV HA constructs $HA_{wt}$-mCherry2 and mCherry2-$HA_{TMD}$ were cloned based on the previously described $HA_{wt}$-mEGFP (5) and mEGFP-$HA_{TMD}$ (5) plasmids. $HA_{wt}$-mEGFP contains the full-length HA protein fused to mEGFP at the (intracellular) C-terminus, whereas in mEGFP -$HA_{TMD}$ a large part of the extracellular domain of HA is replaced by mEGFP. To clone mCherry2-$HA_{TMD}$ and $HA_{wt}$-mCherry2, the mEGFP-$HA_{TMD}$, $HA_{wt}$-mEGFP, and mCherry2-C1 plasmids (5) were digested with AgeI and BsrGI to replace mEGFP with mCherry2. The plasmids PA-mTurqouise, NP-mCherry2 and mEYFP-$HA_{TMD}$ were a kind gift from Andreas Herrmann (Humboldt University Berlin, Germany) (6).

The FPV NA construct was cloned by amplifying NA sequence from pHH21-NA (7), and cloned into pcDNA3.1+ (Thermo Fisher Scientific, Waltham, MA, USA , #V79020) via restriction with NheI and AflII. To clone NA-mCherry2, mCherry2 was amplified from mCherry2-C1, and the obtained insert was ligated into NA-pcDNA3.1+ by digestion with NotI and XbaI. The construct contains full-length NA fused to mCherry2 at the extracellular side. The plasmid mApple-NA was derived from PMT-mApple (a kind gift from Thorsten Wohland, National University of Singapore, Singapore).

#### Virus propagation and titration.

For virus propagation, confluent MDCK II cells were infected with the avian influenza FPV virus mutant 1 (kind gift from Michael Veit, Free University Berlin (8)) at multiplicity of infection (MOI) 0.01 in DMEM with 0.2 % (w/v) Bovine Serum Albumin (BSA; Sigma Aldrich, Taufkirchen, Germany), 2 mM L-glutamine, 100 U/mL penicillin, and 100 µg/mL streptomycin and incubated at 37 °C. After one hour, virus inoculum was removed and the cells were washed twice with Dulbecco's phosphate-buffered saline with $Mg^{2+}$/$Ca^{2+}$ (DPBS+/+; PAN-Biotech, Aidenbach, Germany). Fresh infection medium with 0.1 µg/mL TPCK-treated trypsin (Sigma Aldrich, Taufkirchen, Germany) was added to the cells and incubated for 2-3 days at 37 °C. Upon visual observation of a cytopathic effect, the supernatant was harvested and cellular debris was removed by centrifugation (3000 x *g* for 30 min at 4 °C). Virus aliquots were stored at -80 °C.

To measure the plaque-forming units (PFU) of the suspension, MDCK II cells were grown in six-well plates until full confluency was reached. The cells were infected with serial 10-fold dilutions of the virus containing supernatant and incubated for one hour at 37°C. Virus inoculum was then removed and replaced by SeaPlaque agarose overlay medium (1x Minimum Essential Medium (PAN-Biotech, Aidenbach, Germany), 0.9 % (w/v) SeaPlaque Agarose (Biozym Scientific GmbH, Hessisch Oldendorf, Germany), 0.2 % (w/v) BSA, 2 mM L-glutamine, 100 U/mL penicillin, and 100 µg/mL streptomycin). After three days of incubation at 37 °C and 5 % $CO_2$, agarose overlay medium was removed, cells were fixated with 10 % (w/v) formaldehyde (Sigma Aldrich, Taufkirchen, Germany) for one hour and PFU was determined by crystal violet staining (0.05 % (w/v) crystal violet (Sigma Aldrich, Taufkirchen, Germany), 1 % (w/v) formaldehyde, 1 % (v/v) methanol in 1 x PBS) (9).

#### Transfection and virus infection.

Cells were seeded in 35-mm dishes (CellVis, Mountain View, CA, USA) with an optical glass bottom (#1.5 glass, 0.16–0.19 mm) at a density of $6 \times 10^5$ cells per dish. After 24 h, cells were transfected with Turbofect® according to the manufacturer's instructions (Thermo Fisher Scientific, Waltham, MA, USA) by using 200 ng pDNA per dish for the controls or 600 - 1200 ng pDNA per dish for IAV proteins. Briefly, plasmids were incubated for 20 min with 3 µL Turbofect diluted in 50 µL serum-free medium, and then added dropwise to the cells.

When needed, cells were co-transfected with the reverse genetic plasmid set of FPV excluding segment 7 (encoding M). Instead, M1-mEGFP and M2-untagged were used. This co-transfection procedure is referred to in what follows as “all”.

In some cases, cells were infected with a MOI 5 with IAV FPV mutant 1 in infection medium at 5 h post-transfection, first on ice for 15 min and then at 37°C for 45 min. Samples were then rinsed with DPBS+/+ and typically observed 12 to 16 h after infection.

#### Confocal microscopy system and setup calibration for fluorescence fluctuation spectroscopy.

All fluorescence fluctuation spectroscopy measurements were performed on a Zeiss LSM780 system (Carl Zeiss, Oberkochen, Germany) using a Plan-Apochromat 40×/1.2 Korr DIC M27 water immersion objective and a 32-channel GaAsP detector array. Samples were excited with a 488 nm Argon laser (AlexaFluor®488, mEGFP) and a 561 nm diode laser (mCherry2). For measurements with 488 nm excitation, fluorescence was detected between 499 and 552 nm; for 561 nm excitation, between 570 and 695 nm, after passing through a 488/561 nm dichroic mirror (for two color measurements) or 488 nm dichroic mirror (for one color measurements). All measurements with more than one fluorescent species were recorded sequentially to minimize signal cross-talk. To decrease out-of-focus light, a pinhole with size corresponding to one airy unit (~39 µm) was used. All measurements were performed at room temperature (22 ± 1 °C).

At the beginning of each measurement day, the focal volume was calibrated by performing a series of point FCS measurements with Alexa Fluor® 488 (AF488, Thermo Fischer, Waltham, MA, USA) dissolved in water at 30 nM, at the same laser power, with the same dichroic mirror and pinhole size. Beforehand, the signal was optimized by adjusting the collar ring of the objective and the pinhole position to the maximal count rate for AF488. Then, ten measurements at different locations were taken, each consisting of 15 repetitions of 10 s, and the data were fitted using a three-dimensional diffusion model including a triplet contribution. The structure parameter $S$ (defined as the ratio between the vertical and lateral dimension of the theoretical confocal ellipsoid) was typically around 5 to 9, and the diffusion time $\tau_d$ around 35 to 40 µs. The waist $\omega_0$ was calculated from the measured average diffusion time ($\tau_{d,AF488}$) and previously determined diffusion coefficient $D$ of the used dye at room temperature ($D_{AF488}$= 435 μm$^2$s$^{-1}$) (10), according to the following equation:

$$\omega_0 = \sqrt{4\tau_{d,AF488} D_{AF488}} \quad (1)$$

Typical values were 200–250 nm. All measurements were performed at room temperature.

#### Scanning fluorescence (cross-) correlation spectroscopy.

sFCS and sFCCS were used to probe slow diffusive dynamics in lipid membranes as previously described (5, 11-15) with few modifications. Briefly, a line scan of 256 × 1 pixels (pixel size 80 nm) was performed perpendicular to the membrane with 945.45 µs scan time. Typically, 250,000 lines were acquired (total scan time 3 min) in photon counting mode alternating two different excitation wavelengths. Laser powers were adjusted to keep photobleaching below 20 %. Typical values were ~4.7 µW (488 nm) and ~10 µW (561 nm). Scanning data were exported as TIFF files, imported and analyzed in MATLAB (The MathWorks, Natick, MA, USA) using custom-written code. The analysis started with an alignment of all lines as kymographs and then a division into blocks of 1000 lines. In each block, lines were summed up column-wise and the position along the line with maximum fluorescence was determined. This position defines the membrane localization in each block and is used to align all lines to a common origin. Then, all aligned line scans were averaged over time and fitted with a Gaussian function summed to a sigmoidal function (modelling intra-cellular background signal). The pixels corresponding to the membrane were defined as pixels within ± 2.5σ of the peak. To clearly identify the signal originating from the PM, we restricted our analysis to cells in which the surface concentration of the analyzed FP was > 100 monomers/µm². In each line, these pixels were integrated, providing the membrane fluorescence time series *F(t)*. In order to correct for depletion due to photobleaching, the fluorescence time series was fitted with a two-component exponential function and a correction was applied (16). Then, auto-correlation functions (ACFs; g= green channel, r =red channel), and cross-correlation function (CCF) were calculated as follows, using a multiple tau algorithm:

$$G_i(\tau) = \frac{<\delta F_i(t)\delta F_i(t+\tau)>}{<F_i(t)>^2}, \quad (2)$$

$$G_{cross}(\tau) = \frac{<\delta F_g(t)\delta F_r(t+\tau)>}{<F_g(t)><F_r(t)>}, \quad (3)$$

where $\delta F_i = F_i(t) - < F_i(t) >$ and $i = g, r$.

To avoid artefacts caused by long-term instabilities or single bright events, CFs were calculated segment-wise (20 segments) and then averaged. Segments showing clear distortions (typically less than 25% of all segments) were manually removed (12). Furthermore, a model for two-dimensional diffusion in the membrane and a Gaussian confocal volume geometry was fitted to the ACFs and CCF (15):

$$G(\tau) = \frac{1}{N}\left(1 + \frac{\tau}{\tau_d}\right)^{-1/2}\left(1 + \frac{1}{S^2\tau_d}\right)^{-1/2}. \quad (4)$$

Here, the particle number $N$ and diffusion time $\tau_d$ were obtained from the fit. Moreover, diffusion coefficients were calculated using the calibrated waist $\omega_0$ of the focal volume, $D = \omega_0^2/4\tau_d$. The apparent molecular brightness $\varepsilon$ was calculated by dividing the mean count rate detected for each species *i*, $< F_i(t) >$, by the particle number $N_i$ determined from the fit:

$$\varepsilon_i = \frac{<F_i(t)>}{N_i}. \quad (5)$$

Relative cross-correlation values were calculated from the amplitudes of ACFs and CCFs:

$$rel.cc = max\left\{\frac{G_{cross}(0)}{G_g(0)}, \frac{G_{cross}(0)}{G_r(0)}\right\}, \quad (6)$$

where $G_{cross}(0)$ is the amplitude of the CCF and $G_i(0)$ is the amplitude of the ACF in the *i*-th channel (g = green, r = red) (12).

To analyze concentration-dependent oligomerization, the surface concentration was calculated according to the following equations:

$$N_{monomer,protein} = \frac{<I_{protein}>}{<\varepsilon_{monomer,i}>}, \quad (7)$$

$$surface\ concentration = \frac{N_{monomer,protein}}{p_{f,i}\pi\omega_0^2 S}, \quad (8)$$

where $< I_{protein} >$ is the average fluorescence intensity of the protein of interest within a single cell measurement, $\varepsilon_{monomer,i}$ is the average molecular brightness of the monomer control for the corresponding fluorescence species *i*, and $p_{f,i}$ is the probability factor described below in "Brightness calibration and fluorophore maturation". For all SF(C)CS experiments, $p_f$ values in Eq. 8 have been set to 0.7 and 0.6 for mEGFP and mCherry2 respectively, as previously determined (5). By using the effective detection area ($A_{eff} = \pi\omega_0^2 S$), the surface concentration for a protein of interest in expressed in monomeric units per µm$^2$.

#### (Cross-Correlation) Number and Brightness.

(cc)N&B experiments were performed as previously described (5, 12, 13, 18, 19) with few modifications. Briefly, an image stack was acquired over time at a fixed position in the sample, typically consisting of 100 frames. Images of 128 x 512 pixels were acquired by using a pixel size of 70 nm, and 6.3 µs dwell time and alternating two different excitation wavelengths. Laser powers were maintained low enough to keep bleaching below 20 % of the initial fluorescence signal (typically ∼3 µW for 488 nm, and ∼5 µW for 561 nm). CZI image output files were imported into MATLAB using the Bioformats package (20), and analyzed using a self-written MATLAB script implementing the equations from Digman et al. (21) for the specific case of photon-counting detectors, thus obtaining the molecular brightness and number as a function of pixel position. Before further analysis, pixels corresponding to regions of interest (ROI) were selected manually in an image map. To clearly identify the signal originating from the PM, we restricted our analysis to cells in which the surface concentration of the analyzed FP was > 100 monomers/µm$^2$. Next, to correct for lateral drift during the acquisition, frames were aligned to the first frame by maximizing the spatial correlation between sub-selections in consecutive frames, averaged over both channels, as a function of arbitrary translations (22). Finally, brightness-intensity maps were obtained (see e.g. Figure 2 B). These maps show the pixel brightness with a color code in units of counts/dwell time/molecule. The average fluorescence count rate (counts/dwell time) is represented as pixel intensity.

Corrections for bleaching, minor cell movements and specific detector response were performed as described in (12, 19). The calculation of the multimerization state from apparent brightness values is described below in the paragraph "Brightness calibration and fluorophore maturation".

Finally, for two color measurements, the cross variance $\sigma_{cc}^2 = \langle (I_g - \langle I_g \rangle)(I_r - \langle I_r \rangle) \rangle$ was calculated for each pixel (18). In order to obtain a ccN&B analogue of Eq. 6, we defined:

$$N_{cc} = \frac{\sigma_{cc}^2}{\varepsilon_g \varepsilon_r}, \qquad (9)$$

where $\varepsilon_i$ is the channel brightness ($i = g, r$) calculated as usual as $\frac{\sigma_i^2}{\langle I_i \rangle} - 1$.

The relative cross-correlation values were calculated in analogy to Eq. 6 from the particle numbers for each channel $N_i$ ($i = g, r$; $N_i = \frac{\langle I_i \rangle}{\varepsilon_i}$), and the apparent number of complexes $N_{cc}$:

$$rel.cc. = max\left\{\frac{N_{cc}}{N_g}, \frac{N_{cc}}{N_r}\right\}. \qquad (10)$$

To analyze concentration-dependent oligomerization, the surface concentration was calculated according Eqs. 7 and 8, using $p_f$ values determined daily.

#### Brightness calibration and fluorophore maturation

The molecular brightness, i.e. the photon count rate per molecule, is used as a measure for the oligomeric state of protein complexes. This quantity is often based on the assumption that all fluorophores within an oligomer are fluorescent. However, FPs can undergo dark state transitions or be in a non-mature, non-fluorescent state (23). To quantify the amount of non-fluorescent FPs, we consider all these processes together in a single parameter, the apparent fluorescence probability ($p_f$), i.e. the probability of a FP to emit a fluorescence signal. We used the median of the normalized FP homo-dimer brightness $\varepsilon_{dimer}$ to determine the probability $p_f$ for each FP species *i*:

$$p_{f,i} = \frac{\langle \varepsilon_{i,dimer} \rangle}{\langle \varepsilon_{i,monomer} \rangle} - 1. \qquad (11)$$

(11)

An estimate of the oligomeric state is determined by normalizing the molecular brightness $\varepsilon_i$ by the average molecular brightness $\varepsilon_{i,mono}$ of the corresponding monomeric reference and, subsequently, using the previously determined values of $p_{f,i}$ for species *i* (5):

$$\text{Oligomerization} = \frac{1}{p_{f,i}}\left(\frac{\varepsilon_i}{\langle \varepsilon_{i,monomer} \rangle} - 1\right) + 1. \qquad (12)$$

We applied this transformation to every brightness data point of both (cc)N&B and sF(C)CS measurements, obtaining then the true oligomeric size of the complexes. The $p_f$ was determined daily.

#### Calibration of bi-directional plasmids

We examined bi-directional plasmids with either i) mp-mEGFP upstream and mp-mCherry2 downstream of the bi-directional promoter region (mp-mEGFP ↔ mp-mCherry2) or ii) mp-mCherry2 upstream and mp-mEGFP downstream of the bi-directional promoter region (mp-mCherry2 ↔ mp-mEGFP). sFCS measurements were independently performed at the membrane of transfected HEK293T cells (Figure S1 A) to calculate the concentrations of each FP, for each bidirectional plasmid. The box plot (Figure S1 B) with the single data points for each experiment shows the relative expression ratio, defined as the ratio between the measured amounts of FPs (downstream and upstream of the promoter region). Each FP amount is quantified as the number of molecules detected in the confocal volume, provided by sFCCS. The average of the measured relative expression ratios (4.16) was then used to estimate the concentration of M2 for the experiments in which the M2 ↔ mCherry2 construct was used for transfection. For the sake of simplicity, all three proteins are assumed to mature with similar efficiency.

#### Quantification of the plasmid composition

Cells were co-transfected with six different fluorescent proteins constructs (PA-mTurquoise, M1-mEGFP, mEYFP-HATMD, NA-mApple, NP-mCherry2, and mCardinal-M2). For the acquisition of reference spectra, cells were transfected with only one plasmid. Imaging was performed on a Zeiss LSM780 system (Carl Zeiss, Oberkochen, Germany) using a 40x, 1.2NA water immersion objective operating in the "lambda mode". Samples were excited with a 405 nm laser (mTurquoise), 488 nm Argon laser (mEGFP, mEYFP) and a 561 nm diode laser (mCardinal, mCherry2, mApple). To split excitation and emission light, 405/505 nm and 488/561 nm dichroic mirrors were used. Fluorescence was detected in spectral channels of 8.9 nm (26 channels between 459 nm and 690 nm) on a 32 channel GaAsP array detector operating in photon counting mode. Afterwards, a reference spectrum from each individual FP was defined by using the ''Automatic Component Extraction (ACE)'' function of the

''Linear Unmixing'' module of the ZEN software. These reference spectra were then used to separate the fluorophore channels in cells expressing all six plasmids.

#### Quantification of the PM convexity

Cells were transfected with (i) mp-mEGFP ("control") or M1-mEGFP together with either (ii) mCherry2-M2 ("M1-M2") or (iii) the recombinant virus plasmid set without M1 ("M1-all"). A fourth sample consisted of cells transfected with M1-mEGFP and, 4 h post-transfection, infected with FPV ("M1-FPV"). Cells were imaged after 16 h and analyzed with ImageJ (http://imagej.nih.gov/ij/). First, we generated a binary mask of the cell: the fluorescence images were subjected to Gaussian filtering (radius = 1 pixel) and segmented by applying Otsu threshold (Image → Adjust → Threshold). Next, single outlier pixels were removed (Process → Noise → Remove Outliers), the inner regions of cells were filled (Process → Binary → Fill Holes) and outlines of the final objects were generated (Process → Binary → Outline). The binary outline masks were thus used for the following shape descriptive analysis. We calculated the convexity (also called roughness) of the cells by using the ImageJ plugin "shape-filter" (https://imagej.net/plugins/shape-filter). This parameter measures local irregularities in contour shapes by comparing the perimeter of the cell´s convex hull enclosing the cell to the perimeter of the cell itself:

$$convexity = \frac{perimeter\ of\ convex\ hull}{perimeter}. \tag{13}$$

A value significantly below one indicates the presence of an irregular boundary.

#### Supplemental Figures

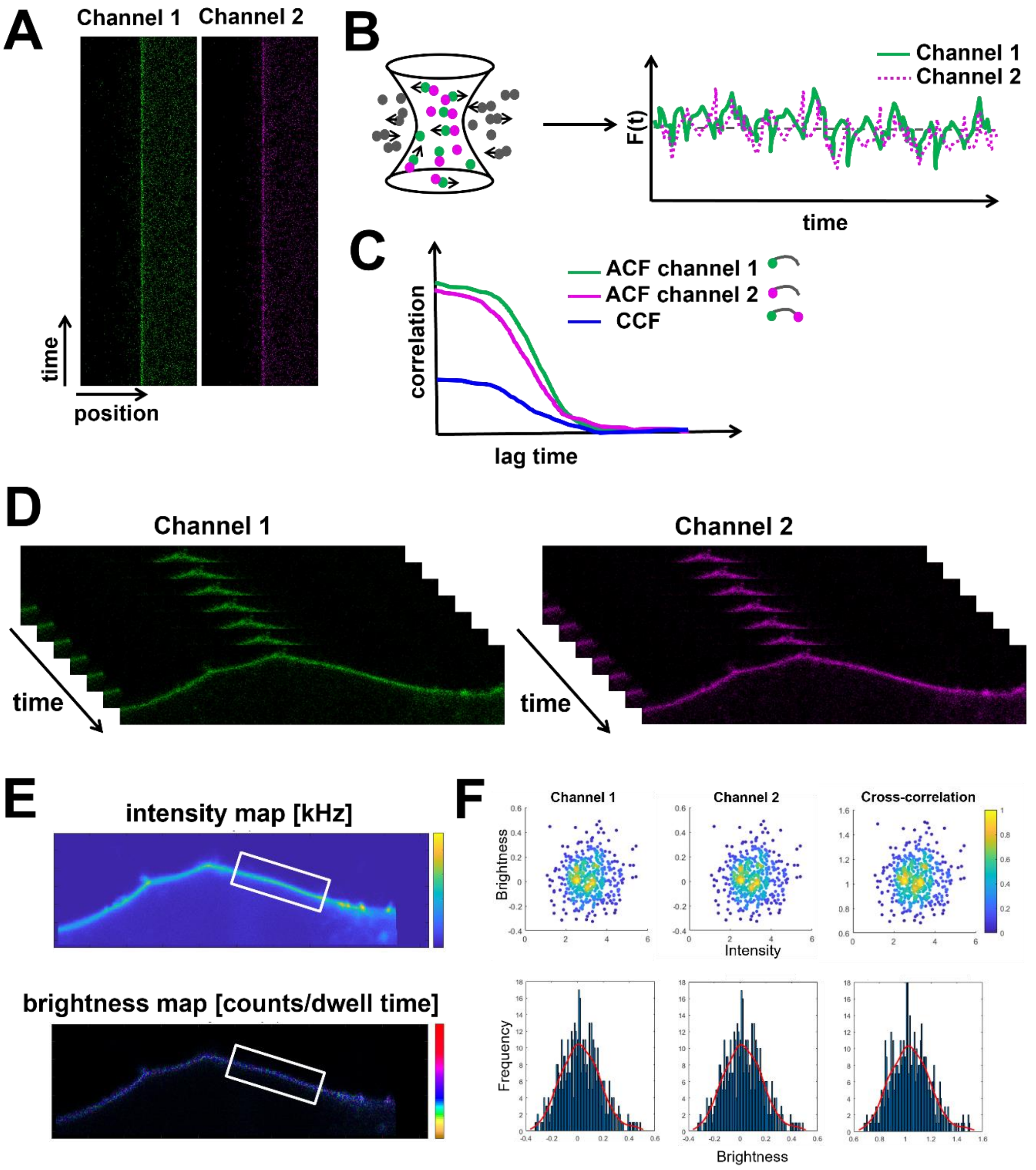


**Figure S1: Schematic representation of scanning fluorescence cross-correlation spectroscopy (sFCCS) and cross-correlation Number and Brightness analysis (ccN&B) in cells.** A: sFCCS measurements are performed by scanning a line perpendicular to the cell membrane in two spectral channels (channel 1, green and channel 2, magenta). Scan lines (represented here as kymographs) are aligned, and membrane pixels are summed. B: sFCCS provides the average number and transit time of fluorescent molecules diffusing through the confocal observation volume. C: Auto-correlation functions (ACFs) of each fluorophore species are calculated from the time-intensity trace, $F_i(t)$ (B), and are represented here in green and magenta. The interaction between two different types of molecules diffusing through the observation volume is determined by calculating the cross-correlation function (CCF) between the two intensity traces. The CCF is represented in blue. D: ccN&B acquisition results in a three-dimensional (x-y-time) image stack. E: Intensity maps and brightness maps of the image stack are obtained from moment analysis of the image stacks and are used to define a region of interest (ROI, white rectangle) around the cell membrane. F: Channel and cross-correlation brightness (ε1, ε2, and Bcc) values are calculated and represented for each pixel. The results are then visualized as scatter plot (brightness as a function of intensity) and as histograms, pooling all selected pixels.

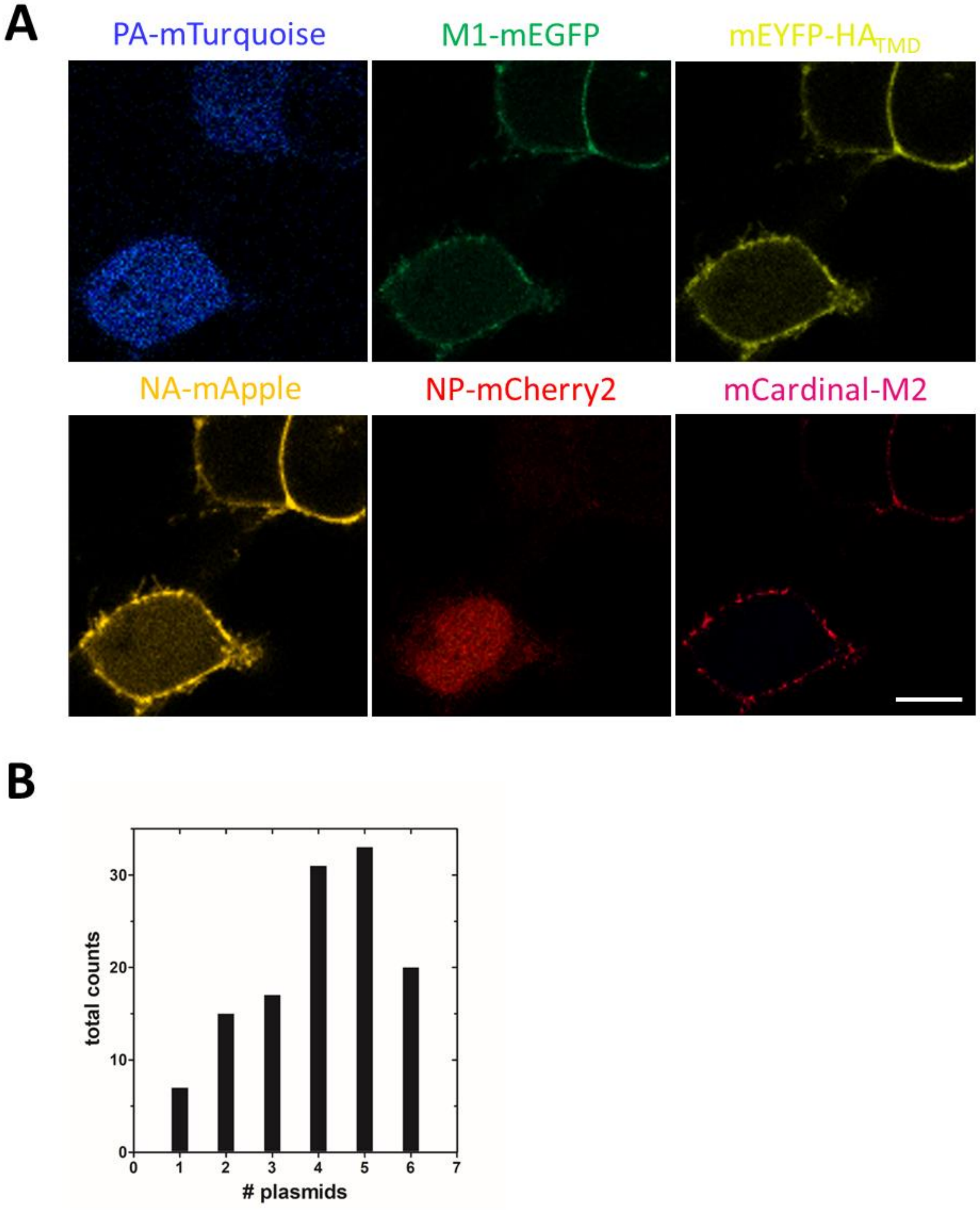


**Figure S2: Quantification of the plasmid expression for six different fusion proteins.** HEK293T cells were transfected with PA-mTurquoise, M1-mEGFP, mEYFP-HA$_{TMD}$, NA-mApple, NP-mCherry2, and mCardinal-M2. After 16 h, samples were imaged using spectral decomposition with channels of 8.9 nm width (26 channels between 459 nm and 690 nm) on a 32 channel GaAsP array detector operating in photon counting mode. A: Representative confocal fluorescence images of HEK293T cells expressing six fusion proteins. Scale bar: 10 µm. B: Quantification of the plasmid expression of each fusion protein in co-transfected HEK293T cells (n = 123). Transfection of the 6 plasmids was performed as described in Material and Methods and in the SI paragraph **"**Quantification of the plasmid composition". The bar plot shows the frequency of the distinct fluorescent proteins observed per cell. According to our results, the majority of visible (i.e. fluorescent) cells expressed > 4 fluorescent constructs (B). The probability to find a fluorescent cell transfected with only one plasmid was circa 5%. The probability of finding a fluorescent cell expressing only M1-mEGFP in these conditions is circa 5%/6=0.8%.

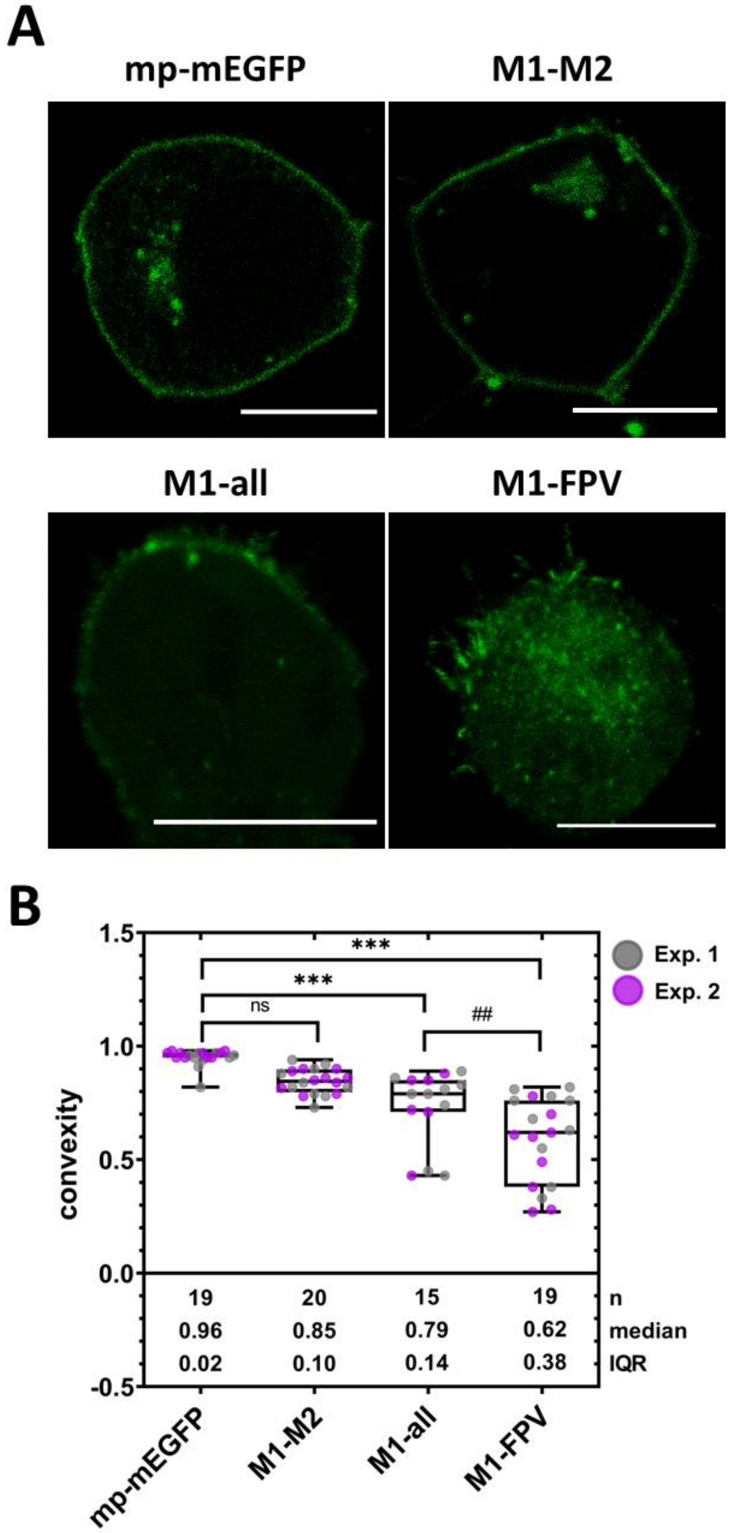


**Figure S3: Shape descriptive analysis of transfected/infected HEK293T cells.** A: Representative confocal fluorescence images of HEK293T cells expressing (i) mp-mEGFP, (ii) M1-mEGFP and unlabeled M2, (iii) M1-mEGFP and the recombinant virus plasmid set (except for segment M, "all") and unlabeled-M2, (iv) M1-mEGFP infected with FPV. A total of 73 images were used for the shape descriptive analysis as described in the SI paragraph "Quantification of the PM convexity". Scale bar: 10 µm. B: Boxplot with single data points (each representing a cell) from two independent experiments shows the convexity of the cell shapes. Cell convexity was calculated by using the ImageJ plugin "shape-filter". Sample size, median, and IQR are indicated in the graph. Statistical significance was determined using one-way ANOVA multiple comparison test (ns indicates not significant, *** indicates $p < 0.001$ compared to mp-mEGFP; ## indicates $p < 0.01$ compared to M1-all). The convexity (also called roughness) of each cell was calculated to quantify local irregularities in contour shapes. The convexity values of M1-all (0.79 ± 0.14, median ± IQR, n = 15 cells) and M1-FPV cells (0.62 ± 0.38, median ± IQR, n = 19 cells) were significantly lower than that measured for control cells expressing mp-mEGFP (0.96 ± 0.02, median ± IQR, n = 19 cells). Cells from the M1-M2 sample (0.85 ± 0.10, median ± IQR, n = 20 cells) showed no significant difference compared to the control. Moreover, M1-FPV cells displayed a significant lower convexity compared to M1-all cells. The higher variance observed in M1-all and M1-FPV cells could be attributed to different expression levels of the respective plasmids and different infection stages/health conditions of the examined cells.

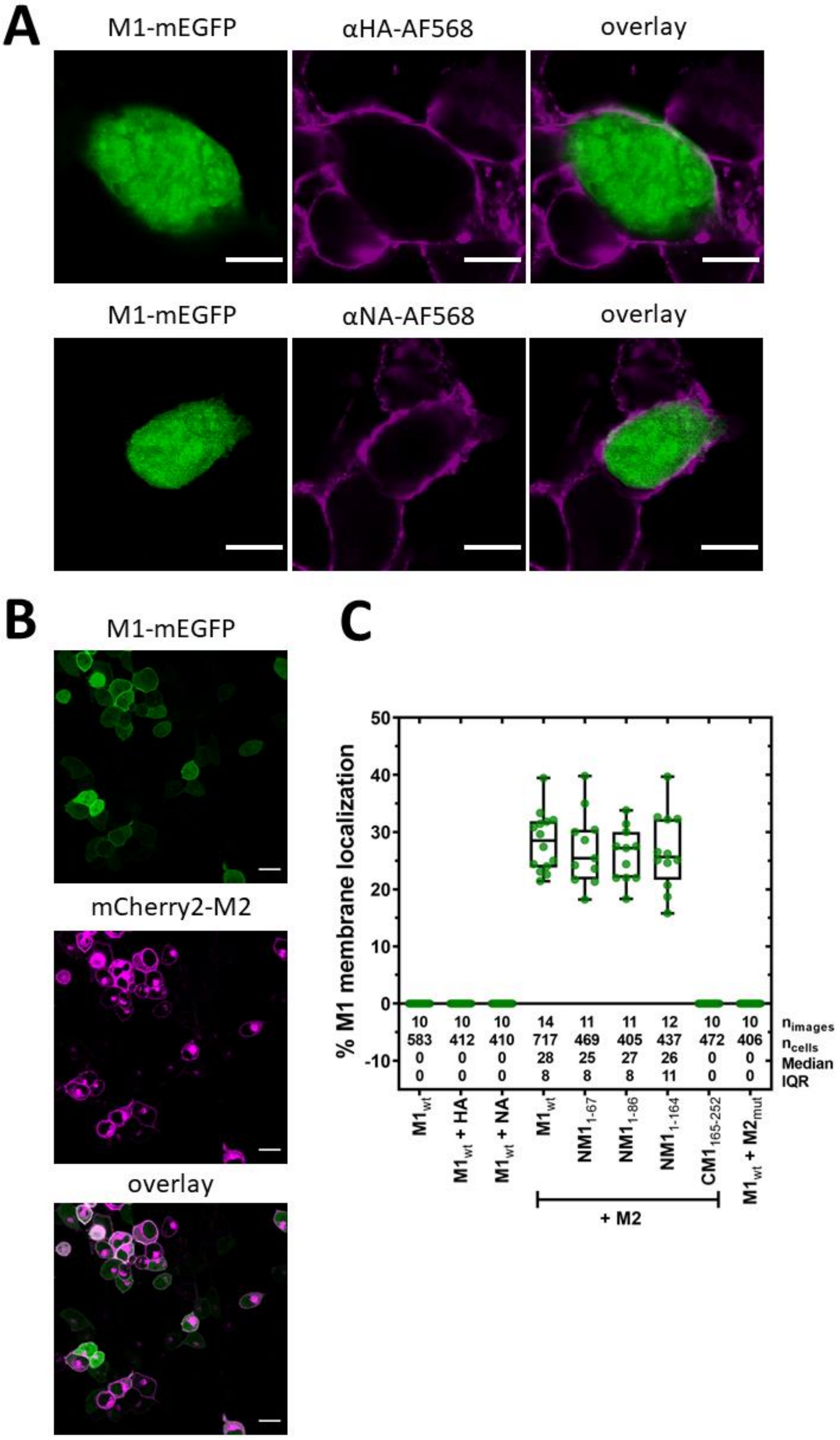


**Figure S4: Immunofluorescence visualization and quantification of membrane-bound M1 in co-transfected cells.** A: Representative confocal fluorescence images of $M1_{wt}$-mEGFP and HA-untagged (upper panel) as well as $M1_{wt}$-mEGFP and NA-untagged (lower panel) co-transfected HEK293T cells after immunofluorescence staining with α-HA-AF568 (magenta) or α-NA-AF568 (magenta). Scale bars are 10 µm. B: Representative confocal fluorescence overview images of $M1_{wt}$-mEGFP and mCherry2-M2 in co-transfected HEK293T cells. Such images were used for the quantification analysis of the recruitment of M1 to the PM. Scale bars represent 10 µm. C: Box Plot with single data points (corresponding to single cells) shows the percentage of HEK293T cells displaying M1 localization at the PM. Sample size, median, and interquartile range (IQR) are indicated in the graph. No recruitment of M1 to the PM was observed in the presence of HA, NA or $M2_{mut}$. In the presence of M2, all the truncated M1-constructs encoding the N-terminal domains were significantly recruited to the PM, whereas the M1 construct encoding only the C-terminal domain showed no membrane localization.

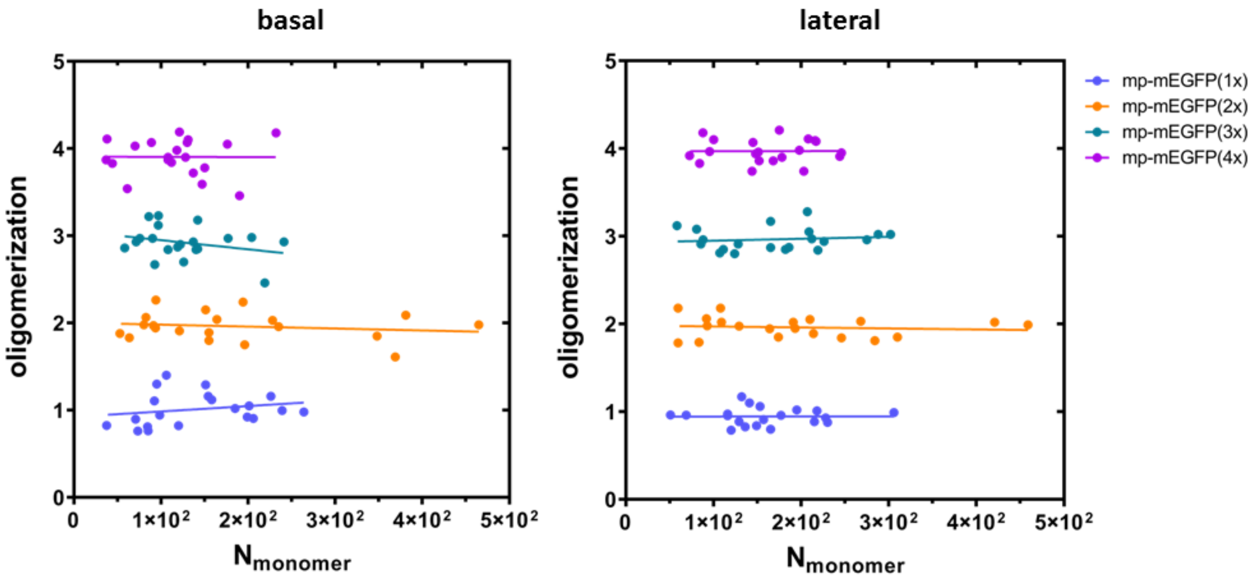


**Figure S5: Comparison of the oligomerization state of oligomeric membrane constructs measured at the basal and lateral membrane in HEK293T cells via N&B.** Oligomerization as a function of the estimated monomer numbers ($N_{monomer}$) for membrane-localized FP constructs (mp-mEGFP(1x) in blue, mp-mEGFP(2x) in yellow, mp-mEGFP(3x) in cyan, mp-mEGFP(4x) in magenta). Values were obtained from N&B measurements at the basal membrane (left panel) and lateral membrane (right panel). Solid lines represent a linear regression fit as guide to the eye. Data are pooled from two independent experiments. The monomer numbers were calculated as ratio between average fluorescence intensity and average monomer brightness. The data show that N&B measurements provide reliable multimerization data for concentrations up to at least ca. $10^3$ monomers / µm² in the described experimental conditions (ca. 5*10² monomers in the confocal volume).

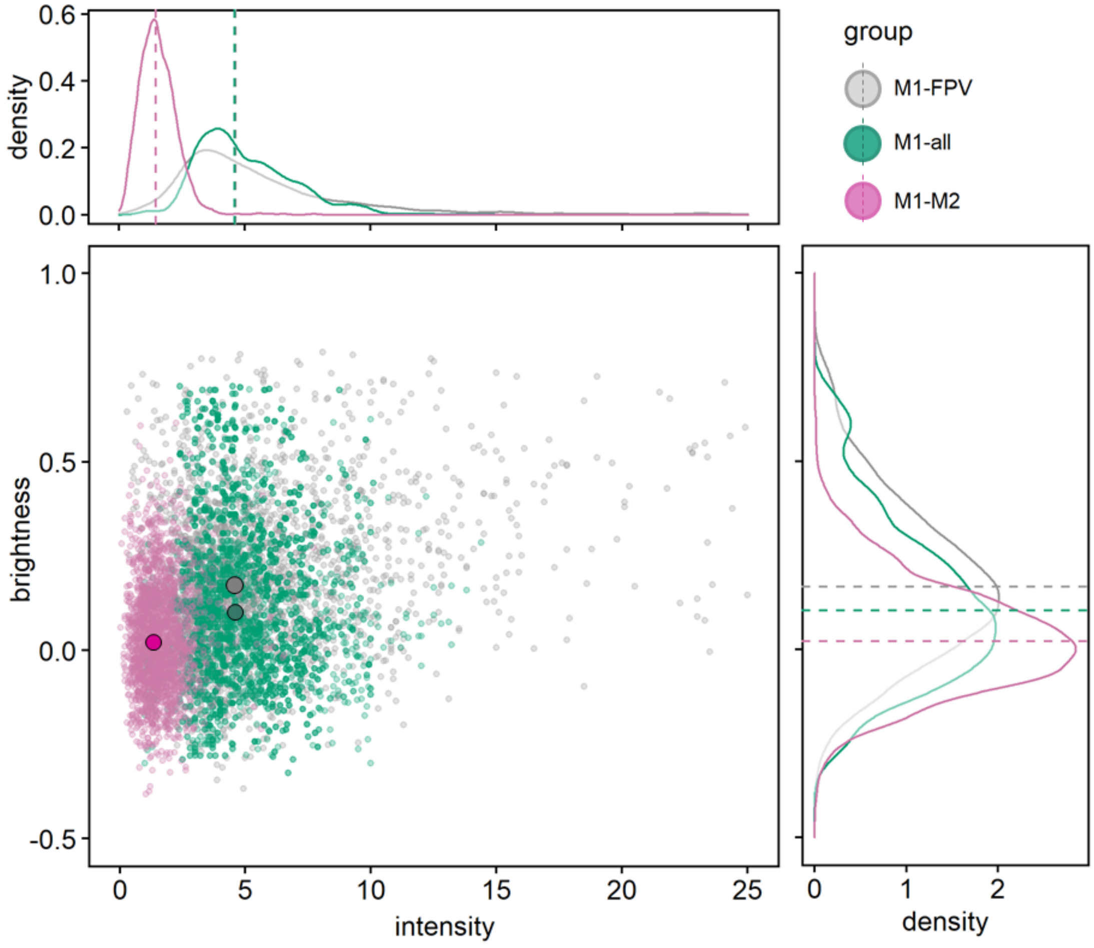


**Figure S6: Higher oligomeric states of M1 in infected cells.** Representative distribution of brightness (counts/dwell time per molecule) and fluorescence intensity (photon counts/dwell time) values for all the pixels within a ROI in an exemplary HEK293T cell expressing i) M1-mEGFP and M2-untagged (magenta), ii) co-transfected cells expressing M1-mEGFP, unlabeled M2 and the reverse genetic plasmid system for all other FPV proteins ("all", green), or iii) M1-mEGFP in FPV infected cells (grey). The brightness-intensity medians are indicated in the graph (large dots with black border line). The sub-panels show the frequency distribution of measured brightness values (right side, solid line), and the frequency distribution of measured intensity values (top side, solid line). The median values of all curves in both sub-panels are shown by the dashed line. Represented data correspond to M1-mEGFP expression at the PM shown in Figure 2 A and B.

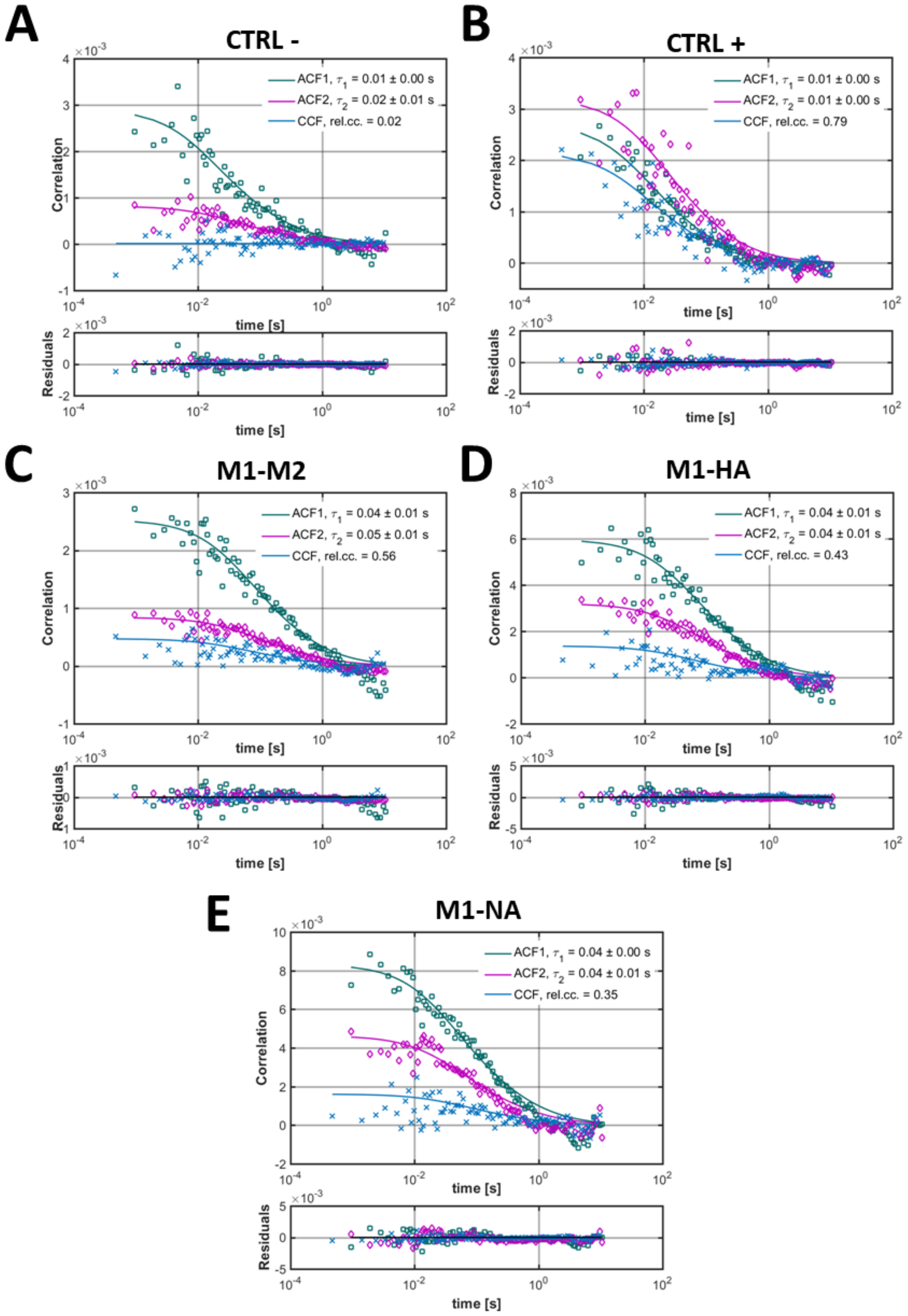


**Figure S7: M1 cross-correlates strongly with M2 and weakly with the glycoproteins HA and NA.** Representative auto-correlation functions (ACFs, mEGFP: green, mCherry2: magenta) and cross-correlation functions (CCFs, blue) obtained from sFCCS measurements on the PM of living HEK 293T cells co-expressing mp-mEGFP/mp-Cherry2 (A, cross-correlation negative control), mp-mCherry2-mEGFP (B, cross-correlation positive control), M1-mEGFP/mCherry2-M2 (C), M1-mEGFP/mCherry2-HA$_{TMD}$ (D), M1-mEGFP/NA-mCherry2 (E). Solid thick lines show fits of a two-dimensional diffusion model to the CFs.

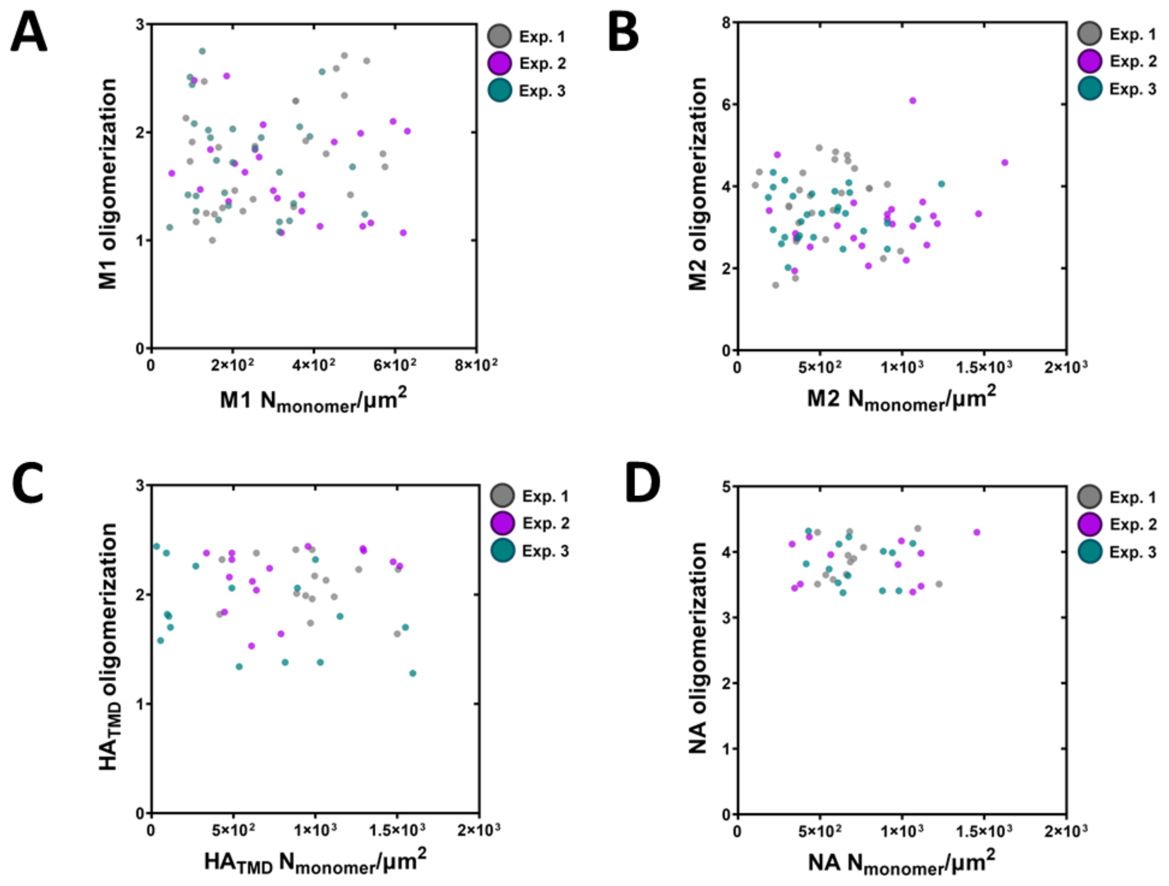


**Figure S8: Oligomerization of viral proteins is independent from surface concentration at the PM.** Scatter plots show oligomerization as a function of surface concentration for M1-mEGFP (A), mCherry2-M2 (B), mCherry2-HA$_{TMD}$ (C), and NA-mCherry2 (D). Values are obtained from sFCCS measurements. Data are pooled from three independent experiments and refer to the data shown Figure 3 of the main manuscript.

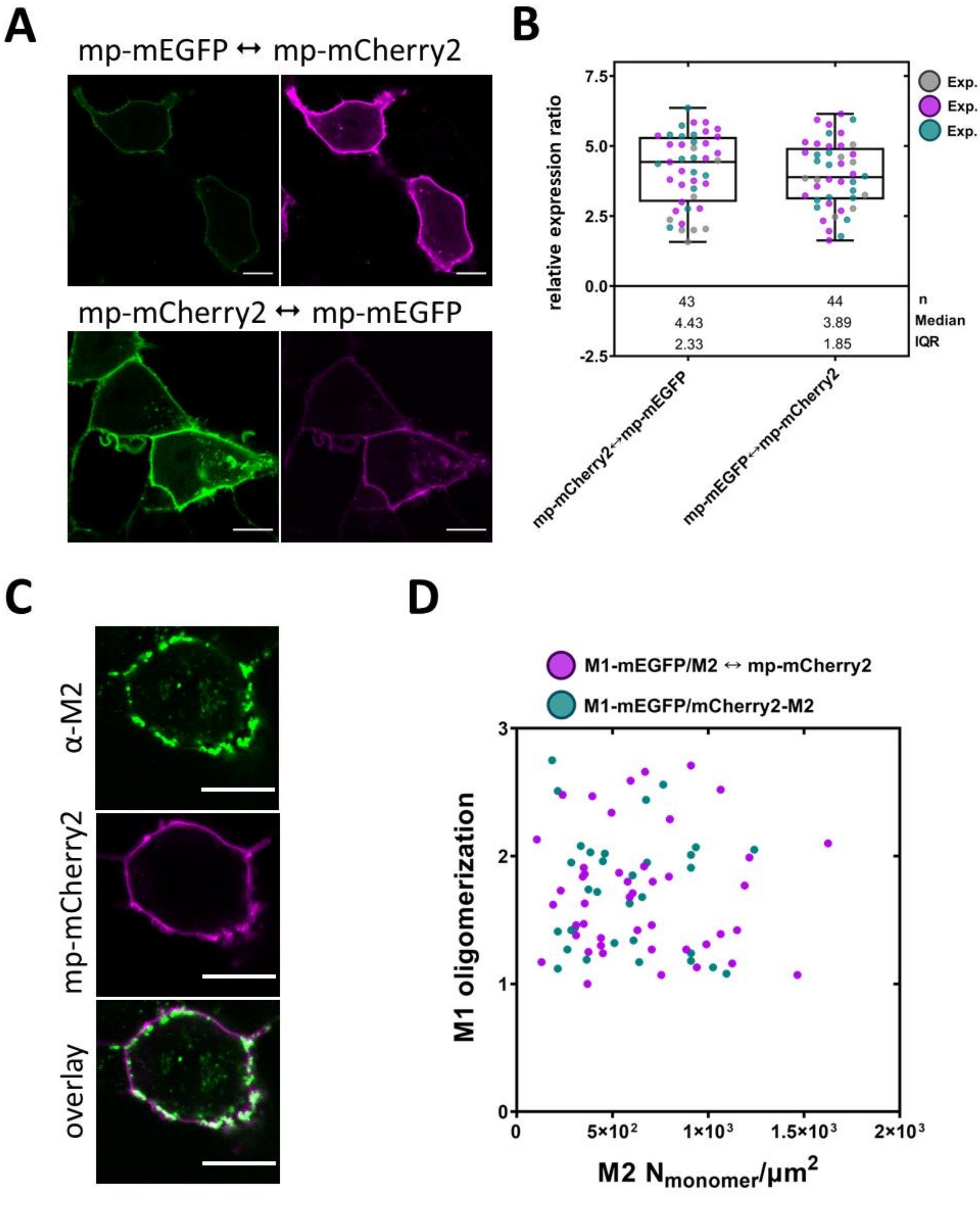


**Figure S9: Expression and calibration of bi-directional plasmids.** A: Representative confocal fluorescence images of the mp-mEGFP ↔ mp-mCherry2 and mp-mCherry2 ↔ mp-mEGFP constructs (mEGFP in green, mCherry2 in magenta) expressed in HEK293T cells. Scale bars represent 10 µm. B: Box plot with single cell data points from three independent experiments shows the relative expression ratios between the surface concentrations of downstream and upstream fluorescent proteins expressed in HEK293T cells. Concentration values were measured using sFCCS at the PM. Sample size, median, and interquartile range (IQR) are indicated in the graph. C: Representative confocal fluorescence images of the M2↔ mp-mCherry2 construct expressed in HEK293T cells after an immunofluorescence staining with α-M2-AF488 (green). Scale bars represent 10 µm. D: M1-mEGFP oligomerization as a function of the estimated M2 surface concentration for the bi-directional M2 ↔ mp-mCherry2 construct, as well as mCherry2-M2. Values are obtained from sFCCS measurements. Data are pooled from three independent experiments.

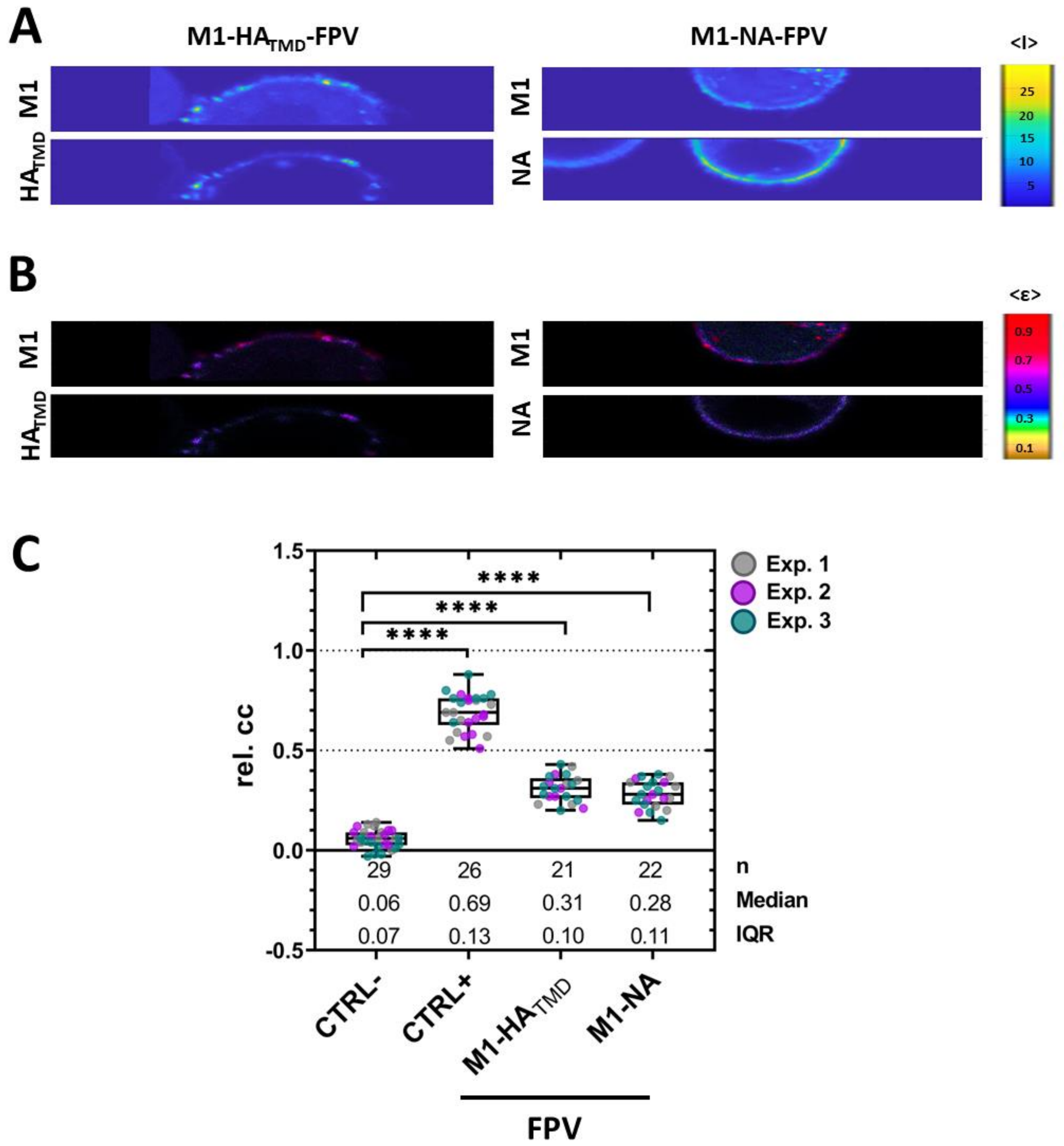


**Figure S10: Weak interaction between M1 and HA$_{TMD}$ or NA in infected cells.** Cross-correlation Number and Brightness (ccN&B) analysis for M1-mEGFP co-transfected with mCherry2-HA$_{TMD}$ or NA-mCherry2 in FPV infected cells. Average intensity as well as molecular brightness maps and rel. cc values were obtained as described in the Methods section. A: Representative average intensity maps of M1-mEGFP (top) in infected HEK293T cells co-expressing mCherry2-HA$_{TMD}$ (bottom, left) or NA-mCherry2 (bottom, right). The average intensity map is visualized via color scale with the unit counts/dwell time. B: representative brightness-intensity maps of M1-mEGFP (top) and mCherry2-HA$_{TMD}$ (bottom, left) or NA-mCherry2 (bottom, right), corresponding to the panels shown in (A). The image shows pixel brightness as pixel color (counts/dwell time/molecule), and mean photon count rate as pixel intensity. C: Box plot with single data points from three independent experiments shows the rel. cc of the controls (negative control: mp-mEGFP(1x)/mp-Cherry2(1x), and positive control: mp-mCherry2-mEGFP), and M1-mEGFP co-expressed with mCherry2-HA$_{TMD}$, and NA-mCherry2. Sample size, median, and IQR are indicated in the graph. Statistical significance was determined using one-way ANOVA multiple comparison test (**** indicates $p < 0.0001$ compared to the negative control (CTRL-)).

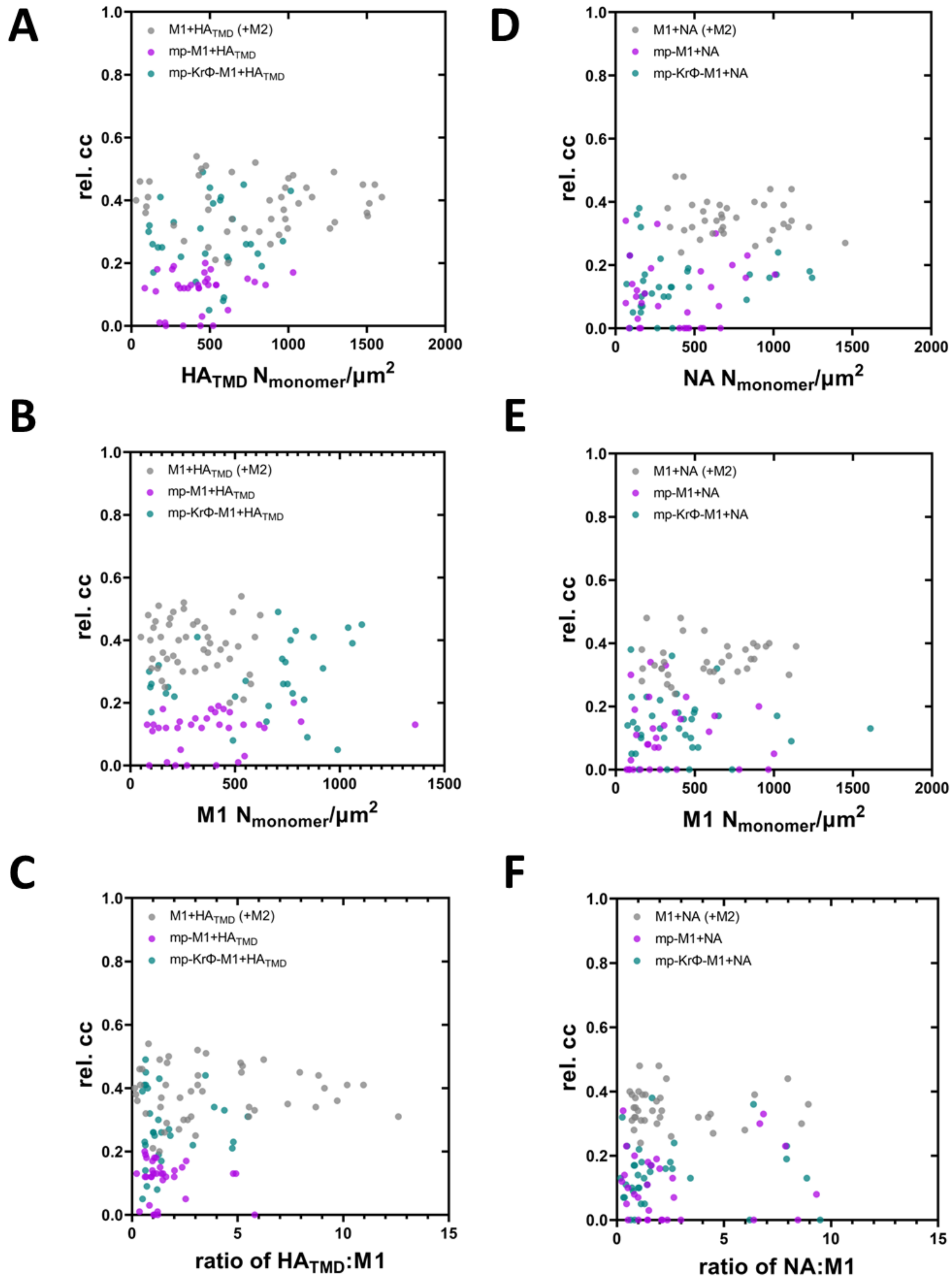


**Figure S11: Cross-correlation between M1 and HA (or NA) is independent from surface concentration at the PM.** A-C: Scatter plots show the rel. cc between different M1 constructs and mCherry2-$HA_{TMD}$, as a function of the surface concentration of mCherry2-$HA_{TMD}$ (A), each of the M1-mEGFP constructs (B), and the expression ratio of mCherry2-$HA_{TMD}$:M1-mEGFP constructs (C). D-F: Scatter plots show the rel. cc between different M1 constructs and NA-mCherry2, as a function of the surface concentration of NA-mCherry2 (D), each of the M1-mEGFP constructs (E), and the expression ratio of NA-mCherry2:M1-mEGFP constructs (F). Solid lines represent a linear regression fit, shown here as guide to the eye. Values are obtained from sFCCS measurements. Data are pooled from three independent experiments and are related to the data shown in Figure 4.
